# Kinesin-1, -2 and -3 motors use family-specific mechanochemical strategies to effectively compete with dynein during bidirectional transport

**DOI:** 10.1101/2022.08.04.502809

**Authors:** Allison M. Gicking, Tzu-Chen Ma, Qingzhou Feng, Rui Jiang, Somayesadat Badieyan, Michael A. Cianfrocco, William O. Hancock

## Abstract

Bidirectional cargo transport in neurons requires competing activity of motors from the kinesin-1, -2 and -3 superfamilies against cytoplasmic dynein-1. Previous studies demonstrated that when kinesin-1 attached to dynein-dynactin-BicD2 (DDB) complex, the tethered motors move slowly with a slight plus-end bias, suggesting kinesin-1 overpowers DDB but DDB generates a substantial hindering load. Compared to kinesin-1, motors from the kinesin-2 and -3 families display a higher sensitivity to load in single-molecule assays and are thus predicted to be overpowered by dynein complexes in cargo transport. To test this prediction, we used a DNA scaffold to pair DDB with members of the kinesin-1, -2 and -3 families to recreate bidirectional transport in vitro, and tracked the motor pairs using two-channel TIRF microscopy. Unexpectedly, we find that when both kinesin and dynein are engaged and stepping on the microtubule, kinesin-1, -2, and -3 motors are able to effectively withstand hindering loads generated by DDB. Stochastic stepping simulations reveal that kinesin-2 and -3 motors compensate for their faster detachment rates under load with faster reattachment kinetics. The similar performance between the three kinesin transport families highlights how motor kinetics play critical roles in balancing forces between kinesin and dynein, and emphasizes the importance of motor regulation by cargo adaptors, regulatory proteins, and the microtubule track for tuning the speed and directionality of cargo transport in cells.

## Introduction

Neurons are elongated, highly polarized cells that require robust, long distance, bidirectional cargo transport to function (Hirokawa & Takemura, 2005). Better understanding of the molecular mechanisms underlying bidirectional cargo transport is needed, as disrupted transport in neurons is linked to neurogenerative diseases including Alzheimer’s, hereditary spastic paraplegia, and amyotrophic lateral sclerosis (ALS) (Bilsland et al., 2010; Chevalier-Larsen & Holzbaur, 2006; De Vos et al., 2008; Gabrych et al., 2019; Millecamps & Julien, 2013; Stokin & Goldstein, 2006; Ström et al., 2008). Intracellular cargo is carried by the cytoskeletal motors kinesin and dynein, which move in opposite directions, toward the plus-end and minus-end of microtubules, respectively (Vale, 2003). Interestingly, it has been shown that kinesin and dynein are simultaneously present on axonal vesicles (Encalada et al., 2011; Hendricks et al., 2010; Maday et al., 2012; Sims & Xie, 2009; Soppina et al., 2009), suggesting that successful bidirectional transport depends on coordination between, and strict regulation of, these antagonistic motors.

The predominant model for bidirectional transport is the “tug-of-war” model, which posits that if both kinesin and dynein are present, they will pull against each other, and the strongest motor will determine the cargo directionality (Gross, 2004). Notable experimental support is the elongation of endosomes immediately preceding a directional switch in *Dictyostelium* cells (Soppina et al., 2009). However, results from a body of experimental and computational studies suggest that the tug-of-war model is not sufficient to account for the range of bidirectional transport activities in cells. Multiple studies have observed a “paradox of codependence” (Hancock, 2014), wherein inhibiting plus-end-directed motors abolishes minus-end-directed movement instead of enhancing it, and vice versa (Gross et al., 2002; Kunwar et al., 2011; Martin et al., 1999). This complexity suggests mechanisms beyond pure mechanical tug-of-war, such as a requirement of both motors for full activation, cargo binding and regulation.

Cellular studies of bidirectional transport have provided important information about the specific isotypes and numbers of motors present on cargo during transport (Cason et al., 2021; Hendricks et al., 2010; Shubeita et al., 2008), as well as characterizing cargo dynamics (Barkus et al., 2008; Kamal et al., 2000; Levi et al., 2006; Maday et al., 2012; Rosa-Ferreira & Munro, 2011; Tanaka et al., 1998), and the codependence of opposite directionality motors (Gross et al., 2002; Kunwar et al., 2011; Martin et al., 1999). However, these studies are limited in their ability to decouple inherent motor properties from external regulation via cargo adaptors, microtubule associated proteins (MAPs), and other factors. *In vitro* optical trap studies have provided precise measurements of the force generation capabilities of single, as well as teams of motors (Andreasson, Shastry, et al., 2015; Budaitis et al., 2021; Gennerich et al., 2007; Guydosh & Block, 2006; Hendricks et al., 2012; Rai et al., 2016; Sanghavi et al., 2021). However, in recent work, traditional single-bead optical trap experiments have been shown to impose non-negligible vertical forces on motors that may accelerate their detachment rate under load (Khataee & Howard, 2019; Pyrpassopoulos et al., 2020). One remedy for this problem is the three-bead trap assay, used widely in studies of myosin (Finer et al., 1994), which significantly minimizes vertical forces and provides a more physiologically relevant measurement of motor behavior under load (Howard & Hancock, 2020; Pyrpassopoulos et al., 2020). Still, in these three-bead traps, the movement of a gliding microtubule is being measured and direct tracking of the motor is difficult. Therefore, assays that precisely measures motor behavior under physiologically relevant loads and without extra confounding variables, are needed.

To directly track kinesin and dynein motor pairs *in vitro*, an elegant method was developed that fuses single-stranded DNA to each motor, links them together through complementary DNA base pairing, and tracks the motor pairs by two-color total internal reflection fluorescence (TIRF) microscopy (Belyy et al., 2016). This method has been used extensively to investigate the mechanical competition between kinesin-1 and various activated dynein complexes bound to BicD2 (DDB), BicDR1 (DDR), and Hook3 (DDH) (Belyy et al., 2016; Elshenawy et al., 2019; Feng et al., 2020; Ferro et al., 2020). These studies report that while DDB can substantially slow down the stepping of kinesin-1, kinesin-1 still dominates kinesin-DDB transport. On the other hand, DDR and DDH, which are more likely to contain two dyneins and may more effectively activate dynein (Grotjahn et al., 2018; Urnavicius et al., 2018), pull kinesin-1 toward the minus-end more often than DDB (Elshenawy et al., 2019). In neurons and other cells, dynein complexes also transport cargo against members of the kinesin-2 (Hendricks et al., 2012; Hendricks et al., 2010; Loubéry et al., 2008) and kinesin-3 (Schuster et al., 2011) families. Importantly, kinesin-2 and −3 families have motility and force generation properties that are distinct from kinesin-1 (Andreasson, Shastry, et al., 2015; Arpag et al., 2019; Budaitis et al., 2021; Chen et al., 2015; Feng et al., 2018; Lessard et al., 2019; Mickolajczyk & Hancock, 2017; Shastry & Hancock, 2010; Zaniewski et al., 2020), which is suggested to play an important role in fast axonal transport. Kinesin-1 can withstand substantial hindering forces for long durations, meaning it is not prone to detaching under load (Blehm et al., 2013; Pyrpassopoulos et al., 2020; Schnitzer et al., 2000; Visscher et al., 1999), but some members of the kinesin-2 and kinesin-3 families have been shown to rapidly detach under load (Andreasson, Shastry, et al., 2015; Arpag et al., 2019; Budaitis et al., 2021). Interestingly, these kinesin-2 and kinesin-3 motors have also been shown to reengage with the microtubule and resume force generation at faster rates than kinesin-1, perhaps compensating for the rapid detachment (Andreasson, Shastry, et al., 2015; Arpag et al., 2019; Budaitis et al., 2021; Feng et al., 2018). Despite these fascinating observations, it remains unclear how the different motile properties of these diverse kinesins affect their coordination with dynein complexes during bidirectional transport.

A recent computational study that used a stochastic stepping model to simulate bidirectional transport found that the different properties of kinesin-1 and kinesin-2 motors substantially affected the directionality and velocity of cargo transport with DDB (Ohashi et al., 2019). There were three key results from this study: i) the magnitude of the stall force for either the kinesin or DDB motors had a negligible effect on the overall cargo velocity, ii) DDB-kinesin-1 pairs had an average cargo velocity near zero while DDB-kinesin-2 pairs had an average cargo velocity of −300 nm/s, and iii) the sensitivity of detachment to load was the strongest determinant of the net cargo velocity. It is notable that despite DDB having a lower stall force parameter than kinesin-2 (4 pN vs 8 pN), DDB-kinesin-2 pairs had primarily DDB-directed cargo motility (average velocity of - 300 nm/s) in the simulations. Overall, these results suggest that force-dependent detachment, rather than stall force, is the best metric for predicting bidirectional transport behavior of a particular set of motor pairs or teams, but this hypothesis needs to be experimentally confirmed.

In the current study, we reconstituted DDB-kinesin bidirectional transport by linking the motors together with complementary single-stranded DNA, and directly tested how the different motility properties of members of the kinesin-1, kinesin-2 and kinesin-3 families impacted the resulting bidirectional motility of DDB-kinesin complexes. Surprisingly, we found that, when analyzing events where both motors were engaged and moving on the microtubule, Kin2 and Kin3 motors were able to withstand hindering loads from DDB nearly as well as Kin1. A stochastic stepping simulation of the three motor pairs support a mechanism by which the fast reattachment kinetics of Kin2 and Kin3 counteracts their rapid detachment under load and enables robust force generation against DDB motors. These results confirm the idea that load-dependent detachment and reattachment are the key parameters that determine motor performance under load and point to family-specific mechanochemical strategies to achieve successful cargo transport.

## Results

### Reconstituting DDB-kinesin bidirectional transport in vitro

To create DDB-kinesin motor pairs, we first expressed and purified constitutively active C-terminal SNAP-tag fusion constructs for well-characterized members of the kinesin-1, −2 and −3 families: *D. melanogaster* KHC/Kin1, *M. musculus* KIF3A/Kin2, and *R. norvegicus* KIF1A/Kin3 (Fig. 1A). The SNAP tags were substoichiometrically functionalized with a 63 bp DNA oligonucleotide and an Alexa-647 dye to achieve a population of dual-labeled kinesin motors (Fig. 1A). The dynein-dynactin-BicD2 (DDB) complex consisted of full-length recombinant dynein expressed in Sf9 cells, purified dynactin from cow brain, and truncated, recombinant BicD2 (25-424) with a C-terminal GFP tag. We determined the concentration of the oligo-labeled kinesin monomers via an SDS-PAGE shift assay (Fig. 1 Supp. 1A, C and E) and then calculated the concentration of labeled dimers using the percent reduction in the unlabeled (unshifted) band intensity of an SDS-PAGE gel after oligo-labeling (Fig. 1 Supp. 1B, D and F). We also confirmed that each labeled motor exhibited velocity and run lengths similar to previously published values for similar constructs (Fig. 1 Supp. 2 and 3) (Feng et al., 2018; Lessard et al., 2019; Mickolajczyk & Hancock, 2017). The functionalized kinesin was linked to the DDB-GFP complex via a GFP nanobody (GBP) (Kubala et al., 2010) functionalized with the complementary 63 bp DNA oligonucleotide (Fig. 1B). Each of the motor pairs were imaged using two-channel TIRF microscopy to simultaneously track both the kinesin and DDB.

**Figure 1:**
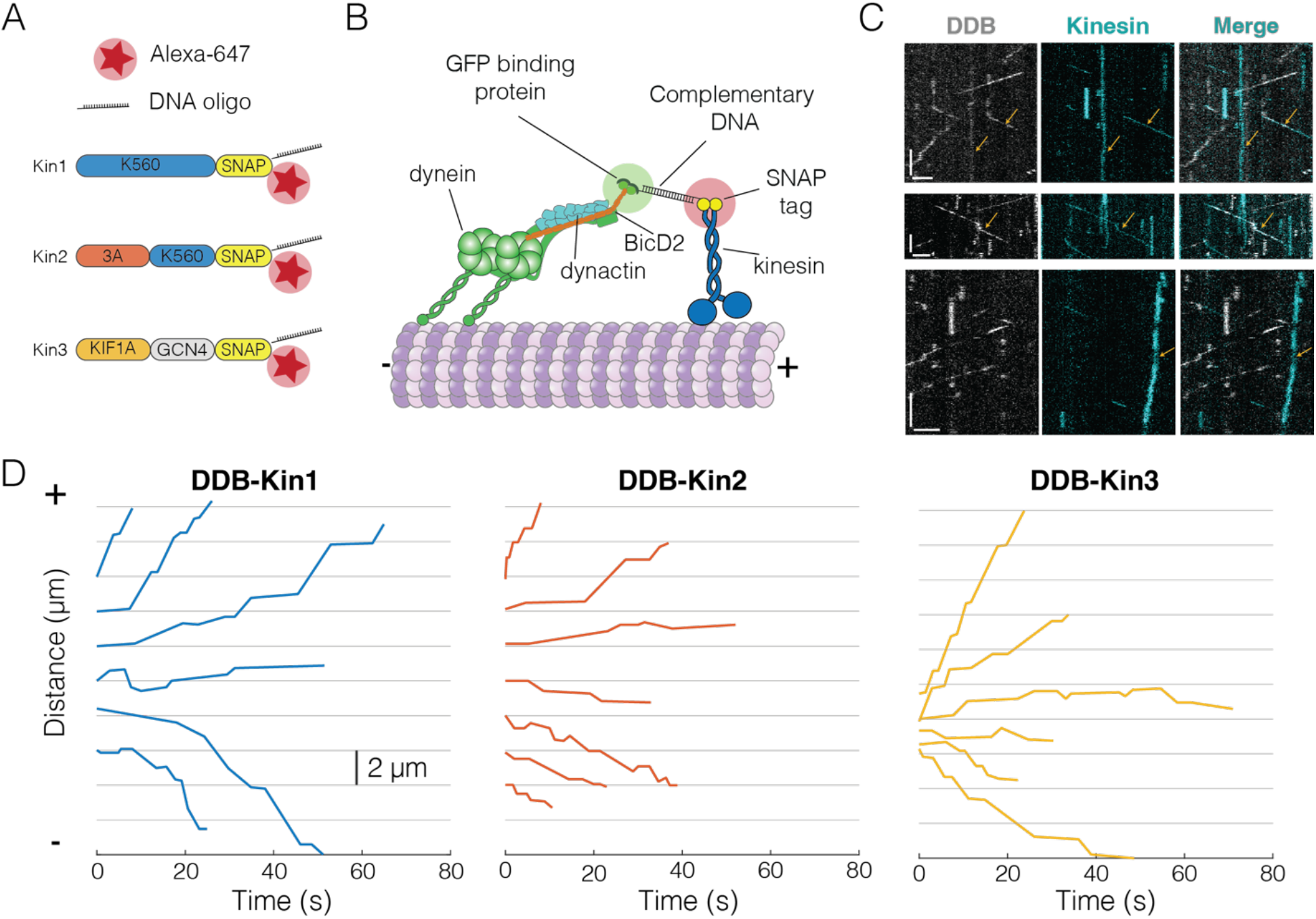
Experimental set-up and visualization of DDB-Kin complexes. (A) Schematic of kinesin constructs containing SNAP tags functionalized with an Alexa-647 dye and a single stranded DNA oligo. (B) DDB and kinesin motors connected via complementary DNA oligos on the GFP binding protein (GBP) and SNAP tag (C) Sample kymograph showing the DDB/GFP channel (gray), the kinesin/Alexa-647 channel (cyan), and the overlay. Scale bars are 2 μm (horizontal) and 10 s (vertical). Microtubule (not shown) is oriented with plus-end to the right. Colocalized events are indicated by an arrow. (D) Sample x-t plots for DDB-Kin1, DDB-Kin2 and DDB-Kin3 complexes. **Figure 1 Source Data uploaded** **Figure 1 Supplement 1:** Purification gels and shift assays **Figure 1 Supplement 2:** Unloaded run length and velocity for Kin1/2/3 **Figure 1 Supplement 3:** Unloaded run length and velocity for DDB **Figure 1 Supplement 4:** Sample traces for DDB-kin1/2/3 pairs

The resulting kymographs included populations of free kinesin motors, free DDB complexes, and colocalized pairs. Microtubule directionality was determined via directionality of the free motors (Fig. 1C). Each set of DDB-Kin traces contained plus-end-directed events with short durations, along with slower events with long durations and net directionality toward either the plus-end or minus-end. Within a single trace, there was considerable velocity heterogeneity, including fast, slow, and paused segments (Fig. 1D, Fig. 1 Supp. 4A-F). Notably, directional switches, defined as sequential segments that move in opposite directions, were rare, occurring with a frequency of 0.01/s for all (Fig. 1 Supp. 4G-I). To understand the differences between the dynamics of the DDB-Kin1, DDB-Kin2, and DDB-Kin3 pairs, we next quantified the overall velocity for each trace and examined differences between the trace velocity distributions for the three motor pairs.

### DDB-Kin1 pairs move toward plus end faster and more often than DDB-Kin2/3 pairs

We first asked how the velocity distributions of the motor pairs compared to velocity distributions of a single kinesin or DDB under zero load. We did this by taking the average velocity of each motor or motor complex over the entire run (trace velocity), where a positive velocity is kinesin dominated motility and a negative velocity is DDB dominated motility. For DDB-Kin1 pairs, the median trace velocity was 290 nm/s compared to 586 nm/s for unloaded kinesin-1; this is a 50% decrease but much faster than has been reported previously (Fig. 2A) (Belyy et al., 2016; Feng et al., 2020). In contrast, the median trace velocities of Kin2 and Kin3 were −28 nm/s and −6.3 nm/s, respectively, which are dramatically slower than the unloaded motor speeds (Fig. 2A). Next, we compared the fraction of complexes that moved with net plus-versus net minus-end directionality. We found that 75% of DDB-kin1 pairs had a net plus-end displacement, whereas only 44% of DDB-Kin2 pairs and 49% of DDB-Kin3 pairs had a net plus-end displacement (Fig. 2B).

**Figure 2:**
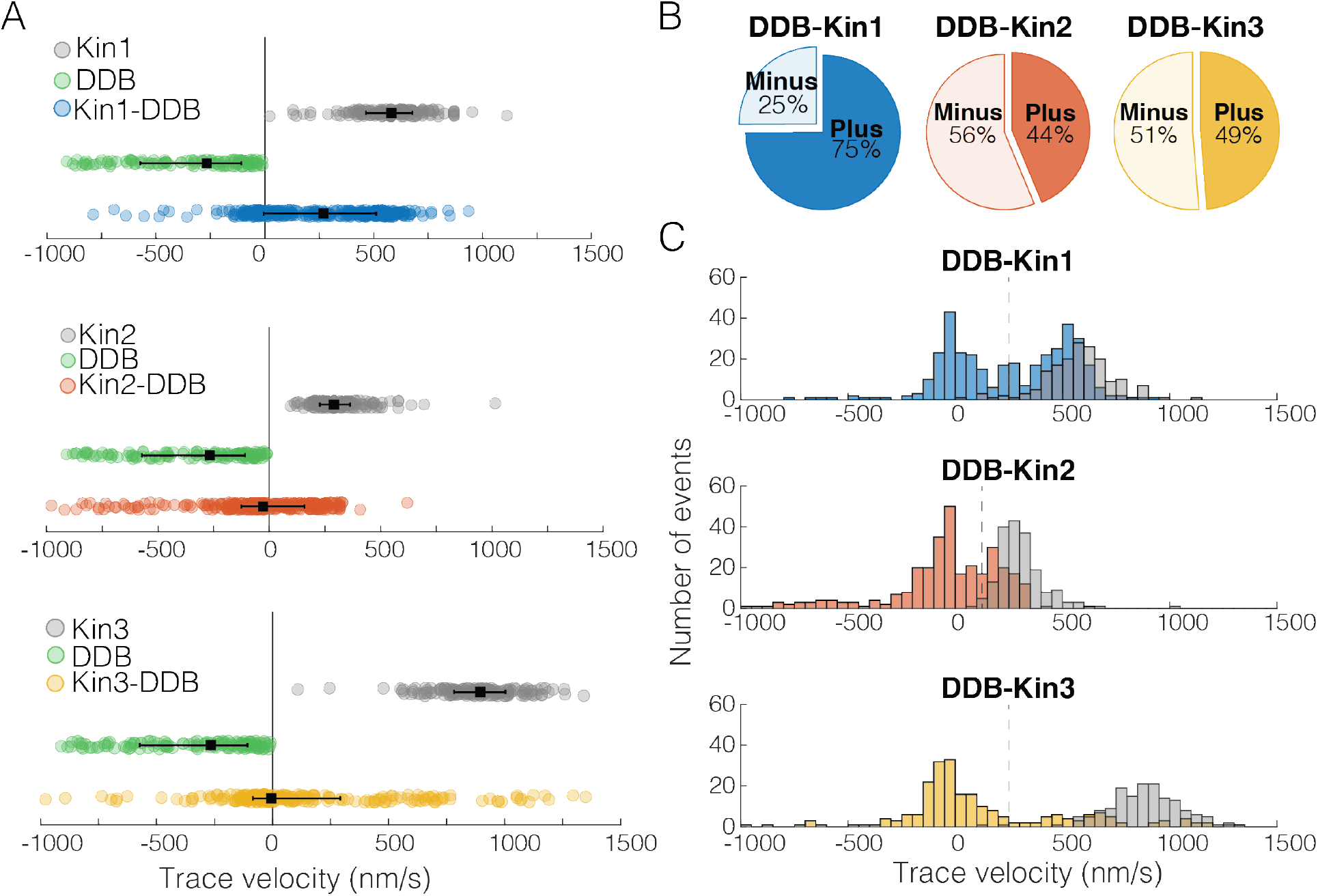
DDB-Kin1 pairs move faster, and more frequently, to the plus-end than DDB-Kin2/3 pairs. (A) Scatter plots showing whole trace velocities of the kinesin alone (gray/top), DDB alone (green/middle) and the DDB-Kin1/2/3 pair (blue/orange/yellow/bottom). Error bars represent median values and quartiles. (B) Fraction of motor pairs having net plus-end displacement (dark blue/orange/yellow) or net minus-end displacement (light blue/orange/yellow) (C) Histogram of the motor pair velocities. Unloaded kinesin velocity distributions are shown in gray (2^nd^ histogram on the right). Dashed line indicates fast-slow trace velocity threshold. **Figure 2 Source Data uploaded**

Notably, the DDB-Kin1 and DDB-Kin2 trace velocity distributions had two clear peaks, one centered near zero and a second centered near the unloaded kinesin motor velocities (Fig. 2C). The DDB-Kin3 velocities had a similar peak near zero, but the fast plus-end population was dispersed rather than centered around a clear peak; this may in part be due simply to the larger range of possible speeds for the faster Kin3. Due to the substantial overlap of the fast plus-end peaks with the velocity distributions for isolated unloaded kinesins, we next investigated whether these two modes represent two configurations of the motor pair: the slow mode representing traces where both kinesin and DDB are engaged on the microtubule and the fast mode representing traces where only the kinesin is engaged.

### Two populations represent only kinesin engaged or both kinesin and DDB engaged on the microtubule

To separate out the fast plus-end population for our analysis, we defined a trace velocity threshold of 250 nm/s for the DDB-Kin1/3 pairs and 125 nm/s for the DDB-Kin2 pairs. We picked these values based on the clear separation of peaks in the trace velocity histograms, and because > 95% of the unloaded velocity data lie above this threshold (Fig. 2C). The fraction of the plus-end events above these velocity thresholds was 52% for DDB-Kin1, 29% for DDB-Kin2, and 25% for DDB-Kin3, suggesting that Kin1 is more likely to pull DDB off the microtubule and move at an unloaded speed than Kin2 or Kin3. However, it is unclear if DDB is fully detached, or remains tethered to the microtubule in a diffusive or weakly-bound state.

To determine whether DDB is detached or in a weakly-bound state, we next compared the motility of the fast motor pairs (pairs with trace velocities above the thresholds) with the motility of unloaded kinesin in single-molecule assays. Comparing run lengths, we found that for Kin1 and Kin2, the presence of DDB caused an ~180% enhancement of the run length (Fig. 3A). In contrast, for Kin3 there was a 47% reduction in the run length in the presence of DDB (Fig. 3A); however, this can be explained by selection bias of long microtubules in the control data, whereas, to maximize the number of events captured in the first few minutes of imaging, DDB-Kin3 events were collected from microtubules of varying lengths. Motor velocities were also affected: coupling with DDB caused a 14% reduction for Kin1 and a ~25% reduction for both Kin2 and Kin3 (Fig. 3B). The fact that the velocities were still relatively fast, but DDB had differential impacts on the different kinesin families, is consistent with DDB being in a diffusive or weakly-bound state that both tethers the kinesin to the microtubule to enhance the run length and creates a frictional drag that has a greater effect on Kin2 and Kin3 motors.

**Figure 3:**
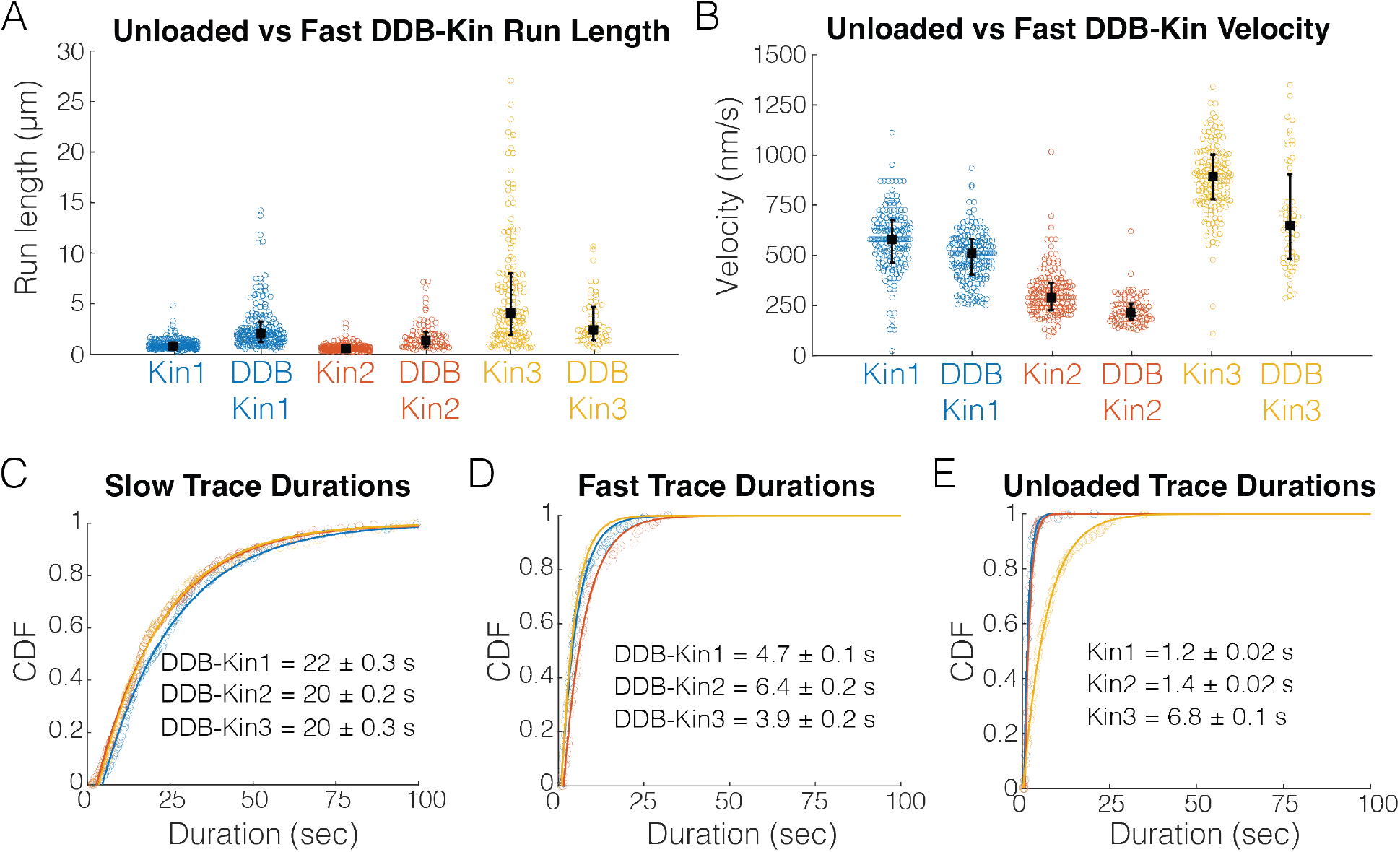
Fast, plus-end events represent a diffusive or weakly-bound DDB. (A) Run length distributions of the fast traces and unloaded kinesin. Error bars represent median values and quartiles. (B) Velocity distributions of the fast DDB-Kin traces and unloaded kinesin. Error bars represent median values and quartiles. (C) Durations of the slow traces (traces < velocity threshold). Data were fit to a single exponential and values are mean duration ± 95% CI of bootstrap distributions. (D) Durations of the fast traces (traces > velocity threshold). Data were fit to a single exponential and values are mean duration ± 95% CI of bootstrap distributions (E) Durations of the unloaded kinesin traces. Data were fit to a single exponential and values are mean duration ± 95% CI of bootstrap distributions **Figure 3 Source Data uploaded**

To better understand the differences between the slow and fast DDB-Kin populations, we next compared the trace durations. The mean durations of the slow traces were 22.2 ± 0.3 sec for DDB-Kin1, 20.3 ± 0.2 sec for DDB-Kin2, and 19.5 ± 0.3 sec DDB-Kin3 (mean ± 95% CI of bootstrap distributions of a single exponential fit; Fig. 3C). These durations are substantially longer than either the durations of the fast DDB-kin populations (Fig. 3D) or the durations of the unloaded motors (Fig. 3E), further supporting the idea that the fast population of DDB-Kin traces represent only the kinesin walking on the microtubule with the DDB being in a diffusive or other weakly-bound state. It follows that the slow velocity, long duration population of DDB-Kin traces represent cases where both motors are engaged in a strongly bound state. Therefore, to explore more deeply how kinesin and dynein connected to a shared cargo compete during bidirectional transport, we focused on the motility of these slow DDB-Kin pairs.

### Pauses observed in motor pairs are due to DDB switching into a “stuck” state

To understand the behavior of the complex when both motors are engaged, we first segmented each trace into segments of constant velocity and plotted the resulting segment velocity distributions (Fig. 4A). Interestingly, there were no clear peaks corresponding to the unloaded DDB or kinesin velocities (Fig. 4 Supp. 1), suggesting the fraction of time where one motor is detached, or not providing a hindering load, is minimal. However, for all three kinesins, the segment velocity distributions had a clear peak at zero velocity, suggesting that one, or both, of the motors spend a significant fraction of time in a static paused state. If these paused states are caused by periods where the kinesin and DDB are pulling with equal force, we would expect the time spent in these paused states to vary between the three kinesin families due to their different propensities to backstep or detach from the microtubule under load (Andreasson, Shastry, et al., 2015; Budaitis et al., 2021; Feng et al., 2018; Ohashi et al., 2019).

**Figure 4:**
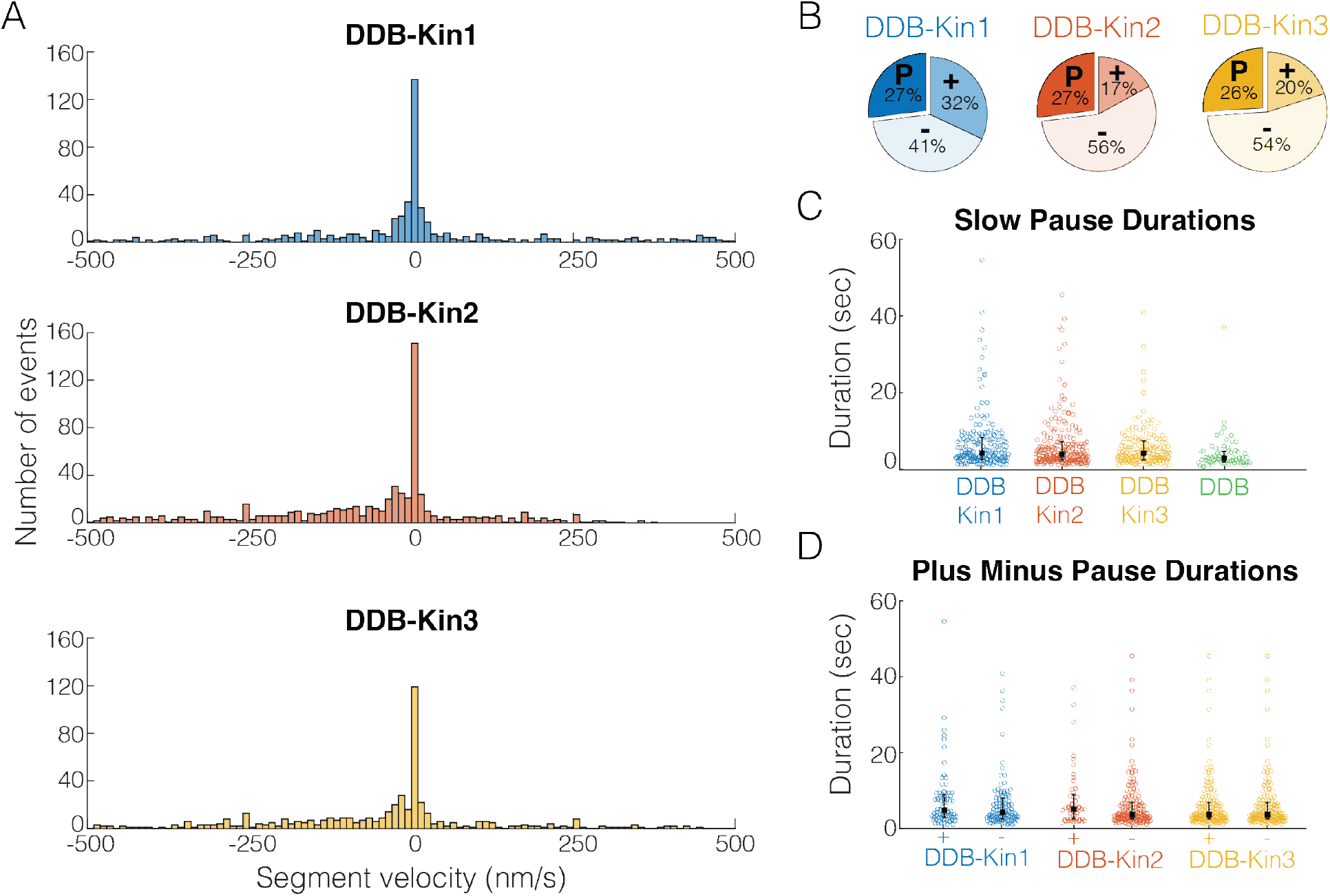
Pauses are due to a DDB “stuck” state. (A) Distributions of segment velocities for the slow traces. <10% of the data is excluded to zoom in on the peak at zero (B) Fraction of segments that are paused (defined as moving < 1 pixel/73 nm), minus-end-directed and plus-end-directed for the slow population (< velocity threshold defined in Fig. 2C). (C) Distributions of pause durations for DDB-Kin1/2/3 pairs compared with unloaded DDB. Error bars represent median values and quartiles. (D) Comparison of pause durations for the minus-end and plus-end-directed events for each motor pair. Error bars represent median values and quartiles. **Figure 4 Source Data uploaded** **Figure 4 Supplement 1:** Slow segment velocity distributions **Figure 4 Supplement 2:** Sample traces for DDB alone

To test whether the fraction of time spent in a paused state was different between the pairs, we quantified the fraction of time that each motor pair spent in each motility state – moving toward the plus end, moving toward the minus end, and paused. A paused segment was defined as any segment that moved less than one pixel (73 nm) in either direction, and plus- and minus-end moving segments involved displacements of more than one pixel. We found that, although the fraction of time spent moving toward the plus-end or minus-end varied across the motor pairs, each motor pair spent a similar 26-27% of the time in a paused state (Fig. 4B). Thus, the fraction of time spent paused is independent of the kinesin type involved and is likely an inherent property of DDB. To confirm the pauses are inherent to the DDB motility, we measured the fraction of time that isolated DDB spends in a paused state and found that it was 24% – almost identical to the motor pairs (Fig. 4 Supp. 2). And in further support of this idea, a previous high-resolution tracking study that rigorously characterized DDB state switching reported that unloaded DDB spends 31% of its time on a microtubule in a “stuck” state (Feng et al., 2020).

To further test whether pulling forces by linked kinesin motors affect the DDB paused state, we next asked whether the duration of the DDB pauses were altered when paired with a kinesin. First, we compared the pause durations of the motor pairs with the pause durations of unloaded DDB and found that compared to the unloaded DDB pause segment durations of 2.8 ± 0.08 sec (mean duration ± 95% CI of bootstrap distributions), the paused segment durations for the motor pairs were 5.1 ± 0.06 sec for DDB-Kin1, 4.5 ± 0.06 sec for DDB-Kin2 and 4.3 ± 0.05 sec for DDB-Kin3 (Fig. 4C). The longer durations indicate that the linked kinesin does not pull DDB out of a paused state, and the ~30-45% enhancement in pause duration suggests that pulling forces from linked kinesins may actually stabilize the DDB paused state somewhat. Further support for kinesin forces elongating the DDB pause state was the finding that pauses that interrupted plus-end events were up to 22% longer than pauses that interrupted minus-end-directed events (Fig. 4D). Based on this slight enhancement in the pause durations between DDB alone and the DDB-Kin pairs, and between plus-end and minus-end-directed events, we conclude the pauses are due to a stabilized version of the DDB “stuck” state rather than a brief stalemate in the tug-of-war.

### Kin1, Kin2, and Kin3 can all effectively withstand DDB hindering loads

Since pause durations suggest that the pauses are due to DDB switching into an inactive static state rather than a property of the mechanical tug-of-war, we decided to focus only on the slow, non-paused segments where both motors are necessarily engaged and stepping along the microtubule. To do this, we separated out the paused segments, and broke the remaining velocity segments into 1-second intervals. The distributions of these 1-second instantaneous velocity intervals were plotted for the DDB-Kin1, DDB-Kin2 and DDB-Kin3 pairs (Fig. 5) to determine if there were any significant differences between the median speed, spread of the distributions, or the fraction of time that complexes move toward the plus-end.

**Figure 5:**
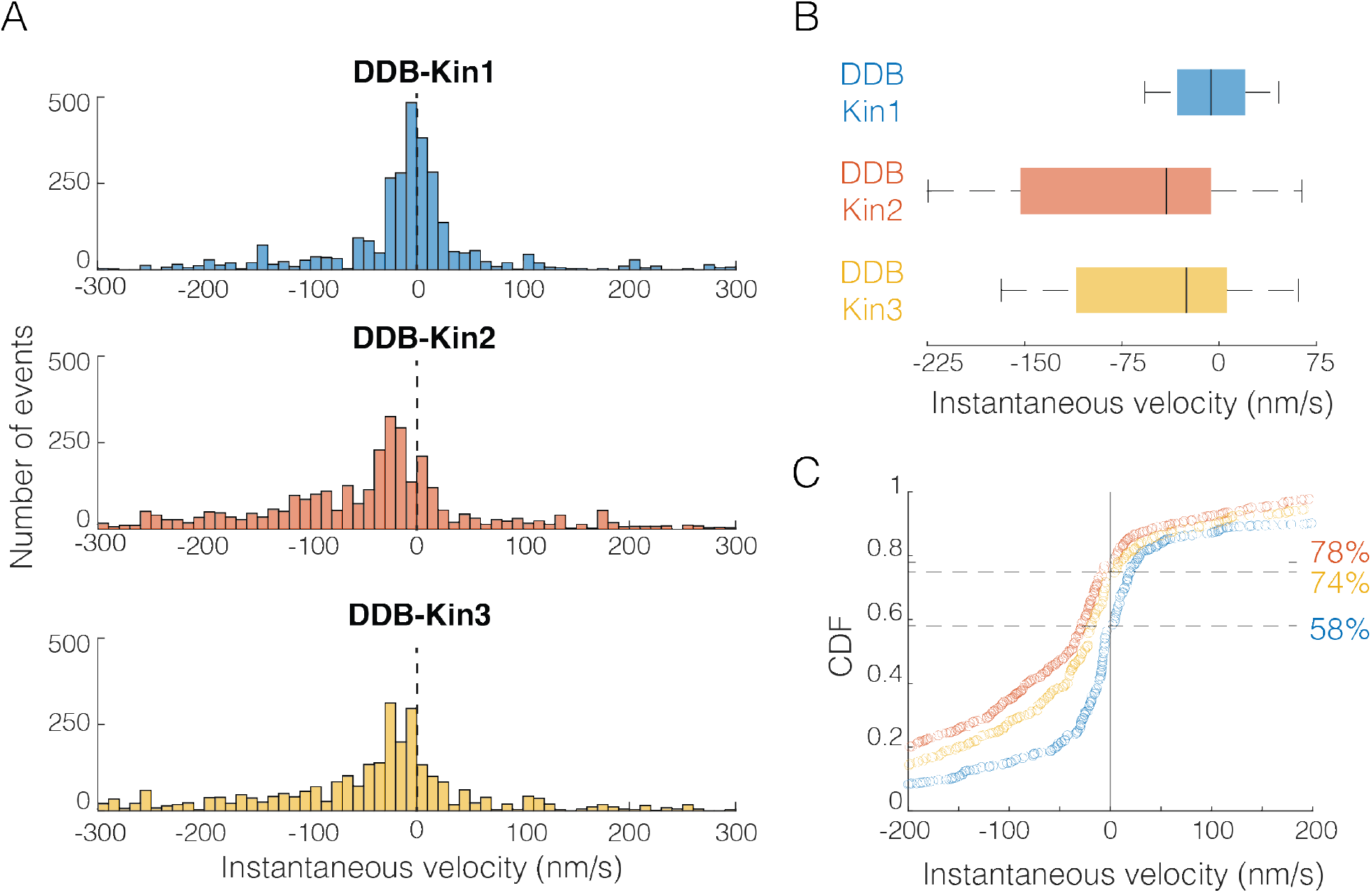
DDB-Kin1/2/3 all compete effectively against DDB._ (A) Distribution of instantaneous velocities calculated over 1-sec time windows for the <u>moving</u> segments (excluding pauses). Dashed line represents v = 0 nm/s. <13% of data are not shown to zoom in on the peak near zero. (B) Box plot distributions of the instantaneous velocity distributions shown in (A). Vertical bars represent median values (−6 nm/s, −41 nm/s, and −26 nm/s), solid boxes represent quartiles, and error bars denote limit of outliers. (C) Cumulative distributions of instantaneous velocities, showing the fraction of time spent moving toward the minus-end (<0 nm/s; denoted by dashed lines) versus the plus-end (>0 nm/s). DDB-Kin1 (blue/bottom), DDB-Kin2 (orange/top) and DDB-Kin3 (yellow/middle). **Figure 5 Source Data uploaded**

From this instantaneous velocity analysis, Kin1, Kin2, and Kin3 motors are all able to withstand hindering loads generated by DDB, but some subtle differences emerge that suggest different underlying mechanisms. First, the peak instantaneous velocity was centered around zero for DDB-Kin1, whereas it was shifted toward the minus-end direction for DDB-Kin2 and DDB-Kin3 (Fig. 5A). Second, the median speed of DDB-Kin1 was −6 nm/s, which is slower than DDB-Kin2 at −41 nm/s and DDB-Kin3 at −26 nm/s (Fig. 5B). Third, the DDB-Kin1 velocity distribution was more confined around zero with the 25% and 75% quartiles spanning 52 nm/s, compared to 159 nm/s and 117 nm/s for DDB-Kin2 and DDB-Kin3, respectively (Fig. 5B). Lastly, the fraction of time spent moving toward the plus-end was ~20% higher for the DDB-Kin1 pairs (Fig. 5C). Together, these data suggest that while all three kinesin motors effectively compete with DDB in tug-of-war, kinesin-1 has a slight advantage. Based on this result, we next wanted to understand how the mechanochemical differences between the three kinesin families lead to their surprisingly similar performances against DDB during bidirectional cargo transport. To do this, we performed stochastic simulations of motor stepping in DDB-Kin pairs.

### Simulations show fast detachment under load can be rescued with fast reattachment

To better understand how the family-specific kinesin motor properties determine the tug-of-war outcomes, we simulated the DDB-kinesin motor pairs using a previously developed stochastic stepping model of bidirectional transport (Ohashi et al., 2019). The model incorporates experimentally determined parameters for kinesin and DDB into a mathematical model that dictates whether each motor will step forward, step backward, detach, or reattach from the microtubule at a given time point (Fig. 6A; see methods) (Ohashi et al., 2019). The kinesin stepping rates and unloaded detachment rates, 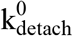, were taken from the single-molecule data in Fig. 3 and Fig. 2 Supp. 1 (summarized in Table 1). The load-dependent motor detachment rate was defined as 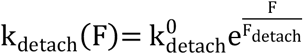, where the force parameter, Fdetach, was taken from published optical tweezer experiments (see Methods) (Andreasson, Milic, et al., 2015; Andreasson, Shastry, et al., 2015; Elshenawy et al., 2019). A number of studies have found that kinesin-3 detaches readily under load (Arpag et al., 2019; Arpağ et al., 2014; Budaitis et al., 2021), and the load-dependent detachment rate for kinesin-3 was taken from a recent study using a three-bead optical trapping assay that minimizes the influence of vertical forces on detachment (Pyrpassopoulos et al., 2022). That same study found that kinesin-3 and, to a lesser extent, kinesin-1 frequently disengage under load, slip backward, and then rapidly reengage with the microtubule. This rapid reengagement behavior has also been observed in other recent studies (Sudhakar et al., 2021; Toleikis et al., 2020). Based on this work, we used reattachment rates of 100/s and 990/s for kinesin-1 and kinesin-3, respectively. Additionally, based on stopped-flow studies that found the microtubule on-rate constant for kinesin-2 is intermediate between that of kinesin-1 and −3 (Feng et al., 2018; Zaniewski et al., 2020), we used a reattachment rate of 300/s for kinesin-2. These fast rates are further supported by the lack of evidence of long periods of kinesin detachment in any previous studies that tracked DDB-Kin pairs (Belyy et al., 2016; Elshenawy et al., 2019; Feng et al., 2020).

**Figure 6:**
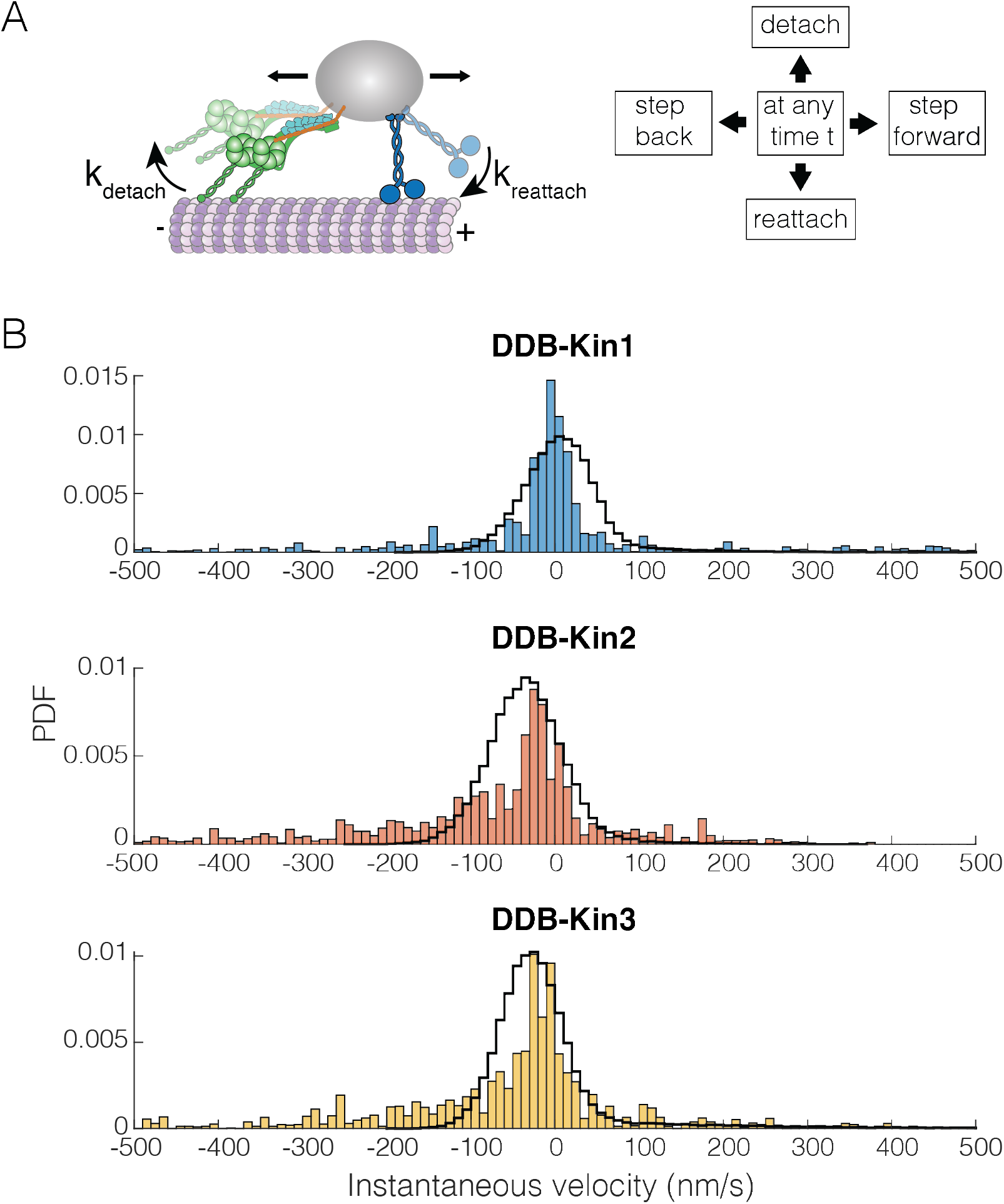
DDB-Kinesin stepping simulations can recapitulate experimental velocities._ (A) Schematic of the stochastic stepping model used in simulations. (B) Instantaneous velocity distributions of the experimental (blue, orange, and yellow bars) and the simulated traces (black lines) for the DDB-Kin1, DDB-Kin2 and DDB-Kin3 pairs. Window size is one second. **Figure 6 Source Data uploaded** **Figure 6 Supplement 1:** Simulation results for slower reattachment rates

**Table 1:** Model parameters.

|  | Kinesin-1 | Kinesin-2 | Kinesin-3 | DDB |
| --- | --- | --- | --- | --- |
| $V_0$ (nm/s) | 586 | 307 | 910 | 360 |
| $k_{\text{forward}} (\text{s}^{-1})$ | 76 | 41 | 117 | 60 |
| $k_{\text{detach}}^0 (\text{s}^{-1})$ | 0.96 | 0.76 | 0.16 | 0.1 |
| $F_s (\text{pN})$ | 6 | 6 | 6 | 3.6 |
| $F_{\text{detach}} (\text{pN})$ | 6.8 | 3 | 1.3 | ∞ (Ideal bond) |
| $k_{\text{reattach}} (\text{s}^{-1})$ | 100 | 300 | 990 | 5 |
| $k_{\text{backstep}} (\text{s}^{-1})$ | 3 | 3 | 3 | 5 |
| $\kappa_{\text{motor}} (\text{pN/nm})$ | 0.2 | 0.2 | 0.2 | 0.2 |

We used the model to simulate bidirectional transport for the three motor pairs and analyzed the resulting trajectories using a 1-second window to calculate instantaneous velocities. Following these initial simulations, we made small adjustments to the parameters to optimize the fits; see Table 1 for final model parameters. Importantly, we found that the model was able to closely match the experimental instantaneous velocity distributions for each case (Fig. 6B and Fig. 6 Supp. 1A). For DDB-Kin1, the peak simulation velocity of 5 nm/s closely matched the experimental peak of −2 nm/s (Fig. 6B). For DDB-Kin2 and DDB-Kin3, the velocity peaks were shifted towards the minus-end compared to DDB-Kin1 and the simulations recapitulated this shift but overshot by ~15-20 nm/s toward the minus-end. One aspect of the experimental data that wasn’t captured by the models was the minus-end tails in the velocity distributions. These tails are likely due to instances of unloaded DDB movement where the kinesin is detached and is slow to rebind because of an unfavorable conformation or geometry, a feature not incorporated into the model.

To examine the role played by motor reattachment kinetics, we repeated the simulations using a reattachment rate of 5 s^-1^ for all three kinesins. This value was initially determined in a study that measured motor-driven deformations of giant unilamellar vesicles (Leduc et al., 2004), it was in a number of modeling studies (Muller et al., 2008), and it was experimentally confirmed in a study that used DNA to connect two kinesins (Feng et al., 2018). When this 5 s^-1^ value was used for the reattachment rate, the simulated velocities had broad distributions centered around −50, −150, and −250 nm/s for DDB-Kin1, 2 and 3, respectively, strongly conflicting with the experimental data (Fig. 6 Supp. 1B). Overall, the data support the conclusion that Kin2 and Kin3 have higher sensitivity to load than Kin1, as seen in the detachment force parameter and in the larger minus-end shift for Kin2 and Kin3 when the slower reattachment rate is used in simulations, but this propensity to detach under load is compensated by fast rebinding to the microtubule. By balancing these attachment and detachment kinetics, all three motors can effectively compete with activated dynein motors during bidirectional transport.

## Discussion

Precise determination of motor behavior under physiologically relevant loads, particularly in the context of bidirectional transport of antagonistic motor pairs, is crucial to understanding how bidirectional transport is regulated in cells (Cason & Holzbaur, 2022; Hancock, 2014). In neurons, members of the kinesin-1, −2 and −3 families are present on cargo alongside dynein (Hendricks et al., 2012; Hendricks et al., 2010; Loubéry et al., 2008; Schuster et al., 2011), but it is unknown why different kinesin motor types are needed and how their differing mechanochemical properties affect their function. Here, we have demonstrated experimentally that Kin1 is only slightly more resistant to detaching under load than Kin2 or Kin3, and all three kinesin motor types generate sufficient force to effectively compete against a dynein-dynactin-BicD2 (DDB) complex. This is seen clearly in Fig. 5 where, although the DDB-Kin2 and DDB-Kin3 peaks are wider and shifted to the left from DDB-Kin1, suggesting more frequent detachment of these kinesin, the median speeds are still close to zero. This result is surprising, as previous computational modeling work found that the strongest determinant of directionality in DDB-kinesin bidirectional transport is the sensitivity of motor detachment to load (Ohashi et al., 2019), and published studies have established that, whereas kinesin-1 is able to maintain stepping against hindering loads, kinesin-2 and kinesin-3 motors detach more readily under load (Andreasson, Shastry, et al., 2015; Arpag et al., 2019; Arpağ et al., 2014; Budaitis et al., 2021; Pyrpassopoulos et al., 2022). Thus, it was expected that kinesin-1 would best compete with DDB, and kinesin-3 would compete the least effectively with DDB.

To understand the unexpected experimental results, we used a stochastic stepping model to simulate the DDB-Kin bidirectional transport. We were able to reproduce the experimental velocity distributions best by implementing fast motor rebinding rates in our stochastic model. Previous work from our lab connected two kinesins using a similar DNA linkage approach to the present work and arrived at a motor reattachment rate of 5 s^-1^ (Feng et al., 2018), a value used widely in published modeling studies (e.g. (Muller et al., 2008)). However, as part of that work, we also made a first principles calculation of the predicted kinesin reattachment rate, 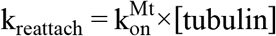. The bimolecular on-rate for microtubule binding in solution, 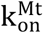, was measured by stopped-flow to be 1.1 μM^-1^ s^-1^ and the effective [tubulin] was calculated to be 125 μM (Feng et al., 2018). Thus, the fast kinesin reattachment rates of 100-1000/s used in the simulations are supported by first principles calculations. The precise values we used in the modeling came from a recent three-bead optical tweezer study that measured the rate that kinesin-1 and kinesin-3 motors reengaged and resumed a force ramp following termination by a rapid backward displacement (Pyrpassopoulos et al., 2022). The large discrepancy between these measured rates of 5 s^-1^ and ~100 s^-1^ could be due to the presence of assisting (Kin-Kin) vs. hindering (DDB-Kin) load, where hindering load may be more likely to optimize fast reattachment due to being pulled back along the microtubule. Importantly, fast reattachment as a strategy to compensate for fast detachment under load explains how seemingly “force-sensitive” kinesin family members can be robust transport motors.

In pairing different types of kinesin motors with DDB, we also gained important insights into the behavior of activated dynein under load. Previous optical tweezer studies have characterized DDB force generation and its stepping behavior under load (Belyy et al., 2016; Elshenawy et al., 2019), and unloaded single molecule assays quantified the switching between processive, diffusive and static states in unloaded single molecule assays (Feng et al., 2020). However, it remained unclear how the switching dynamics would change under load, or how state switching might affect the overall motility when DDB was paired with an antagonistic motor. When full traces were analyzed (Fig 2), there was a sub-population of complexes that moved at the kinesin speeds for each motor pair, which we conclude are most likely due to diffusive or weakly-bound DDB complexes. When we performed a segmental analysis, which focuses on pairs with two active and engaged motors (Fig. 4), we found that DDB state switching kinetics did not change dramatically under load – the DDB-Kin complexes spent ~30% of the time in a static state and the rest of the time in a processive state, similar to DDB alone. The similar duration of the pauses between all three motor pairs suggested that these pauses were due to entirely to DDB switching into a static state, and that this must be a strongly bound state that kinesin cannot pull it out of. Interestingly, the slightly longer duration of these pauses in the DDB-Kin complex suggests that the hindering load provided by the kinesin may actually stabilize this paused state, increasing its duration. Overall, these behaviors suggest that some of the complicated vesicle motility observed in vivo (Hendricks et al., 2010; Rai et al., 2016) could be due to activated dynein switching between states rather than to a tug-of-war between the kinesin and dynein.

Mechanical tug-of-war, where the direction and speed of bidirectional cargo transport is determined by the stronger team of kinesin or dynein motors (Gross, 2004), has been the predominant model for nearly two decades (Hancock, 2014). However, consistent with the present work, recent studies that paired a kinesin with an activated dynein through complementary DNA have observed primarily slow and smooth motility, with no obvious periods of motors moving at unloaded speeds and either zero or very few instances of directional switching (Belyy et al., 2016; Elshenawy et al., 2019; Feng et al., 2020). These results are contradictory to what is predicted by the tug-of-war model (Muller et al., 2008). Interestingly, similar studies of kinesin-dynein bidirectional transport that link the motor pairs with the cargo adaptors TRAK2 and Hook3, rather than DNA, observe primarily fast and unidirectional motility with either zero or very few instances of directional switching (Canty et al., 2021; Fenton et al., 2021; Kendrick et al., 2019). The lack of frequent directional switching in either context suggests that a tug-of-war is not the primary mechanism for determining directionality. Instead, the contrast between the motility of pairs of constitutively active motors linked via DNA versus full-length motors attached to cargo adaptors suggests that cargo adaptors may regulate transport direction by alternately inhibiting kinesin or dynein from engaging with the microtubule, a mechanism termed ‘selective activation’ (Cason & Holzbaur, 2022). Besides providing a more localized site for regulation, direct attachment of antagonistic motor pairs to a common cargo adapter also provides a natural mechanism to ensure that the numbers of plus-end and minus-end motors attached to a cargo are balanced.

An alternative mechanism by which cargo adapters can regulate cargo directionality is by preventing simultaneous binding of kinesin and dynein in the first place. It has been shown that the phosphorylation state (in the case of JIP1; (Fu & Holzbaur, 2013)) or the presence of binding partners (in the case of HAP1; (Twelvetrees et al., 2019; Twelvetrees et al., 2010; Wong & Holzbaur, 2014)) can cause differential binding of kinesin or dynein to a cargo, a mechanism termed ‘selective recruitment’ (Cason & Holzbaur, 2022). Beyond cargo adaptors, the microtubule track itself can also regulate motor binding via recruitment of MAPs that differentially inhibit and recruit motors across the kinesin superfamily, as well as, dynein (Ferro et al., 2022; Monroy et al., 2020). The surprising result that kinesin-1, −2 and −3 can effectively pull an activated DDB during bidirectional transport, despite their drastically different motility characteristics under load, provides strong evidence that motors are not simply regulating themselves via mechanical competition, and underscores the importance of deciphering the combinatorial MAP/adaptor/motor code that regulates transport in cells.

## Acknowledgments

The authors would like to thank Anthony V. Ludlam from the Cianfrocco lab for purifying the dynactin. This work was supported by NIH grants R01GM076476 and R35GM139568 to W.O.H, R21AI152869 to M.A.C., and F32GM137487 to A.M.G.

## Data Availability

Numerical data used to generate figures have been included as source data.

## Code Availability

MATLAB code for the simulations has been uploaded as a source code file for Fig. 6.

## Methods

### Plasmid design

The Kin1 construct consists of *D. melanogaster* KHC residues 1-559 (adapted from Addgene #129761), the Kin2 construct consists of the *M. musculus* KIF3A residues 1-342, followed by the *D. melanogaster* KHC neck coil (345-557) for dimerization (adapted from Addgene #129769), and the Kin3 construct consists of the *R. norvegicus* KIF1A residues 1-393, followed by a GCN4 leucine zipper for dimerization (adapted from Addgene #61665; (Norris et al., 2014)). All of the kinesin constructs have a C-terminal SNAP tag followed by a 6x His tag for purification. The GFP binding protein nanobody, GBP, contains an N-terminal SNAP tag and a C-terminal 6x His tag for purification (Feng et al., 2020; Feng et al., 2018; Kubala et al., 2010). The BicD2 consists of an N-terminal 6x His tag, *M. musculus* BicD2 residues 25-424 (McKenney et al., 2014), followed by GFP, a SNAP tag, and a Strep Tag II.

The dynein plasmid was prepared as described before (Schlager et al., 2014). In summary, the pACEBac1 expression vector containing the dynein heavy chain (DYNC1H1) fused to His-ZZ-TEV-SNAPf tag (pDyn1) and a plasmid containing DYNC1I2, DYNC1LI2, DYNLT1, DYNLL1, and DYNLRB1 (pDyn2) were recombined with Cre recombinase to generate final donor plasmid. pDyn1 and pDyn2 were generous gifts from Andrew Carter, and all genes were codon-optimized for expression in insect cells.

### Protein expression and purification

The kinesin, GBP and BicD2 constructs were bacterially expressed and grown in 800 mL of Terrific Broth (Sigma Aldrich, St. Louis, MO) at 37 °C until the OD = 1-2. Induction was initiated by adding 0.3 mM IPTG and the cultures were left to shake at 21 °C overnight. The cells were harvested and spun at 123,000 x g to collect the supernatant, which was then purified by nickel gravity column chromatography, as described previously (Gicking et al., 2019; Zaniewski et al., 2020). Elution buffer contained 20 mM phosphate buffer, 500 mM sodium chloride, 500 mM imidazole,10 μM ATP and 5 mM DTT. For the BicD2, the final elution peaks were combined, supplemented with 10% glycerol and flash frozen before storage at −80 °C. The concentration was determined using absorbance at 488 nm. The kinesin and GBP constructs were exchanged into 1x PBS with 1 mM DTT and labeled with DNA and SNAP-Surface Alexa Fluor 647 (NEB, Ipswich, MA) dye directly after elution, as described below. Their concentrations were determined using absorbance at 280 nm.

The dynein baculovirus was prepared from pACEBac1 final donor vector using standard methods. High Five™ Cells (BTI-TN-5B1-4) insect cells at 2 × 10^6^/ml density were infected with passage 2 of virus at 1:100 ratio and harvested 72 h later. For 10 ml culture, 1 ml of lysis buffer (50 mM HEPES pH 7.4, 100 mM NaCl, 10% glycerol 10%, 0.5 mM EGTA, 1mM DTT and 0.1 mM ATP, 1 unit Benzonase + SIGMAFAST™ Protease Inhibitor Tablets + 0.5 mM Pefabloc SC) was used. The cell pellet was lysed using a dounce homogenizer at 25 strikes with a tight plunger. Lysate became clear by centrifugation for 88 min at 50K rpm 4°C. Clear lysate was incubated with 0.5 ml packed beads IgG Sepharose 6 Fast-Flow (GE Healthcare, Chicago, IL) for 2-3 hours at 4°C on a tube roller. After incubation, the beads were collected in a disposable column and washed by 150 ml lysis buffer (50 ml +protease inhibitors and 100 ml without protease inhibitors), followed by a wash with 300 ml DynBac TEV buffer (50 mM Tris-HCl pH 8, 2mM Mg-Acetate, 1mM EGTA, 250mM K-Acetate, 10% glycerol, 1mM DTT, 0.1 mM ATP-Mg). The beads were transferred to a 2 ml tube, and 1.5 ml DynBac TEV buffer + TEV protease at final 100 μg/ml was added to the beads. After overnight incubation at 4°C on tube roller, the supernatant was cleared from the beads using low-binding Durapore (0.22 μm). Eluent was subjected to size-exclusion chromatography with Superose 6 300/10 equilibrated with GF150 buffer (25 mM HEPES pH 7.4, 150 mM KCl, 0.5 mM EGTA, 1 mM DTT). The fractions containing dynein were collected and concentrated with 100 KDa MWCO Amicon filter. Glycerol, at final 10%, was added to concentrated dynein before flash-freezing

Dynactin was purified natively from bovine brain following the described procedures (Urnavicius et al., 2015). Fresh cow brains were purchased from the local source and washed immediately with ice-cold PBS at least 3x. With the help of a razor, any significant portions of white matter, blood vessels, membrane, brainstem, and corpus callosum were trimmed away. The collected tissue was washed with PBS 3x again before lysing. 200 ml ice-cold lysis buffer (35 mM PIPES pH 7.2, 1 mM MgSO_4_, 0.1 mM EDTA, 0.2 mM EGTA, 0.2 mM ATP-Mg, 1 mM DTT, Protease inhibitors (cOmplete tablets, or homebrew cocktail + 1 mM PMSF)), 200 μl antifoam, and clean brain tissues were added to a cold metal blender. Tissues were lysed using the “pulse” blender function - 15 seconds on, 15 seconds rest, repeat 4x at 4°C. Lysate was transferred to Oakridge tubes and centrifuged for 1 hr at 15000 g, 4°C. Supernatants from the last step were transferred to Ti45 centrifuge tubes and spun at 40k rpm for 45 min, 4°C. Final clear lysate was loaded into SP-sepharose, 300 ml bed volume, equilibrated with Buffer A (35 mM PIPES pH 7.2, 1 mM MgSO_4_, 0.1 mM EDTA, 0.2 mM EGTA, 0.1 mM ATP-Mg, 1 mM DTT). Bound proteins were fractionated using a two-phase salt gradient: 0% to 25% buffer B (35 mM PIPES pH 7.2, 1 mM MgSO_4_, 0.1 mM EDTA, 0.2 mM EGTA, 0.1 mM ATP-Mg, 1 mM DTT and 1 M KCl) for 900 ml, and then 25% to 100% buffer B for 300 ml. A western blot for p150^Glued^ was used to determine the fractions with dynactin. Fractions with dynactin were collected, diluted twice into HB buffer (35 mM PIPES-KOH pH 7.2, 1 mM MgSO_4_, 0.2 mM EGTA, 0.1 mM EDTA, 1 mM DTT), and loaded into an HB-buffer equilibrated MonoQ 16/10 column. Unbound proteins were washed out with 10 CV of HB buffer, and dynactin was fractionated using three linear gradients: 5% to 15% buffer C (HB buffer + 1M KCl) in 1 CV, 15% to 35% buffer C in 10 CV and 35% to 100% buffer C in 1 CV. Fractions with dynactin were collected and concentrated to 200 μl using 100KDa MWCO Amicon concentrators. Lastly, size-exclusion chromatography of concentrated dynactin (with Suprose6 300/10 column in non-reducing GF150) resulted in a peak of dynactin just after the void volume. At this point the dynactin subunits were distinguishable on 12% SDS-PAGE. Dynactin fractions were concentrated with 100KDa MWCO Amicon concentrators and glycerol was added to the final 10%.

### Functionalizing DNA oligos

Complementary amine-modified 63 bp DNA oligos (IDT) were used. The sequences were /5AmMC12/GT CAA TAA TAC GAT AGA GAT GGC AGA AGG GAG AGG AGT AGT GGA GGT AGA GTC AGG GCG AGA T (kinesin oligo) and /5AmMC12/AT CTC GCC CTG ACT CTA CCT CCA CTA CTC CTC TCC CTT CTG CCA TCT CTA TCG TAT TAT TGA C (GBP oligo). These oligo designs were adapted from previous work (Belyy et al., 2016) and confirmed to have a low probability of forming secondary structures. To functionalize the oligos with BG for SNAP tag binding, 250 μM of each oligo was incubated with 13.28 mM of BG-GLA-NHS (NEB) in 100 mM sodium borate and 50% v/v DMSO. The reaction was then desalted into 1x PBS supplemented with 1 mM DTT using a PD MiniTrap column (Cytiva, Marlborough, MA). The BG-labeling was confirmed using a 10% TBE-Urea electrophoresis gel, and the BG-oligo concentration was determined via absorbance at 260 nm.

### Labeling kinesin and GBP with oligos

BG-oligos were incubated with the SNAP-fusion kinesin and GBP constructs at a 1.5:1 ratio for 1 hr on ice. For the kinesin constructs, 50 μM of SNAP-Surface Alexa Fluor 647 (NEB) was added and incubated for another 30 min on ice to saturate the remaining SNAP-tag binding sites. A second nickel gravity column chromatography purification was performed to separate the labeled protein from the excess BG-oligos and dye. The elution buffer contained 20 mM phosphate buffer, 500 mM sodium chloride, 500 mM imidazole,10 μM ATP and 3-5 mM DTT. The fraction of oligo-labeled monomers was determined by the percent reduction in the unlabeled (unshifted) band intensity on an SDS-PAGE gel, and the concentration of oligo-labeled monomers was determined via an SDS-PAGE shift assay using a gradient of complementary oligo concentrations. The concentration of oligo-labeled dimers was then calculated from the fraction and concentration of oligo-labeled monomers. For the kinesin constructs, the fraction of oligo-labeled monomer was ~50%, which minimizes the fraction of dimers with two oligos on the SNAP tag to < 25%.

For the Kin3 construct, the final oligo-labeled dimer concentration was estimated to be between 1.2 μM – 1.7 μM assuming 20% - 80% oligo-labeling of the monomer. We used the average of 1.4 μM.

### MT Pelleting Assay

To prepare the motors for imaging, a microtubule pelleting assay was performed to remove inactive motors and remove free GBP after incubation with the oligo-labeled kinesin motors. Unlabeled microtubules were polymerized for 30 min at 37 °C in BRB80 supplemented with 1 mM GTP, 1 mM MgCl2 and 10% v/v DMSO. The polymerized microtubules were then diluted in Pelleting Buffer (BRB80 with 100 μM AMPPNP, 10 μM Taxol, 0.3 mg/ml BSA and 0.8 mg/ml casein) to a final concentration of 1.5 μM. The oligo-labeled kinesin were incubated with oligo-labeled GBP on ice for ~10 min and then added to the diluted microtubules at a concentration of 150-300 nM. The microtubule-motor mixture was incubated at room temp (21 °C) for 10 min and airfuged at 25 psi for 10 min to pellet the microtubules. The pellet was resuspended in Resuspension Buffer (30 mM Hepes, 50 mM potassium acetate, 2 mM magnesium acetate, 1 mM EGTA and 10% glycerol, supplemented with 10 μM Taxol, 50 μM ATP, 0.3 mg/ml BSA, 1 mg/ml casein, 0.2 mM glucose, and 0.2 mM βME), incubated at room temp for another 10 min and airfuged at 25 psi for 10 min. The supernatant with the active Kinesin-GBP motors was collected and used for the TIRF experiments. The final concentration of active motors was determined by measuring the 647 fluorescence of the pelleting supe (Unbound Motor), resuspension supe (Active Motor) and the resuspension pellet (Rigor Motor) and using the following formula:

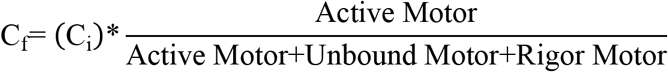

### TIRF assays and analysis

Single-molecule tracking was performed on a custom built micromirror TIRF microscope with a dual-view for two-channel imaging (Nong et al., 2021). All experiments were performed at 21 °C. Unlabeled microtubules were polymerized using the same process described above. Flow cells were prepared by first flowing in 2 mg/ml casein, followed by full-length rigor kinesin (Mickolajczyk et al., 2015). Next, Taxol-stabilized, unlabeled microtubules were flowed in and incubated for 30 sec, unbound microtubule were washed out, and the process was repeated 2x. DDB complexes were formed by combining dynein, dynactin, and BicD2 at a 1:1.5:1 ratio (dynein:dynactin:BicD2), and incubating on ice for 10 min. Kin-DDB pairs linked by complementary oligonucleotides were formed by diluting Kinesin-GBP and DDB complexes to 1-2 nM and incubating them together at an equimolar ratio on ice. Kin-DDB complexes were then introduced into the flow cell in the presence of ~5 μM ATP, allowed to incubate for 2 min, and this process was repeated 1-2x. This low ATP approach maximizes the number of Kin-DDB bound to the microtubules, while retaining activity of the motors. To initiate motility, imaging solution was introduced, consisting of 30 mM Hepes, 50 mM potassium acetate, 2 mM magnesium acetate, 1 mM EGTA and 10% glycerol, supplemented with 2 mg/ml casein, 20 mM glucose, 37 mM βME, glucose oxidase, catalase, 10 μM Taxol, and 2 mM ATP.

Images were taken at 3.5 fps for 100 seconds using a Teledyne Photometrics (Tucson, AZ) Prime 95B sCMOS camera. The two channels from the Dual View split screen were aligned using TetraSpeck microspheres (Invitrogen, Waltham, MA). The composite kymographs were then analyzed manually using Fiji (Schindelin et al., 2012). Consistent directionality of at least three free kinesin and/or DDB motors were required to determine the microtubule polarity. A trace was determined to be a motor pair event if it could be clearly detected in both channels or if it was a single color moving in the wrong direction (e.g. Alexa647/kinesin moving toward the minus end). Whole trace velocities and run lengths were determined by measuring the distance and time over which the moving complex could be observed. All three motor pairs were prepared and imaged on the same day to control for DDB activity, experiments were repeated on three different days to confirm the trend and then the kymographs were pooled together for further analysis. Velocity segmentation of full traces was done manually, where each segment had to be at least 3 frames (858 ms) to be counted, and the minimum detectable velocity change between segments was ± 10 nm/s. Pauses were defined as any segments that moved less than one pixel (73 nm). Directional switches were defined to be sequential segments that moved at least one pixel in opposite directions. Instantaneous velocity distributions were obtained by weighting each velocity or pause segment by its duration (rounded to the nearest second) and plotting the resulting 1-sec segments as a histogram. All of the analysis and plotting was done in MATLAB.

### Simulations

The simulation was a Gillespie stochastic stepping model using an algorithm that followed a previously published model (Ohashi et al., 2019). The reactions of each motor in the simulation were forward stepping, backward stepping, detaching from the microtubule and reattaching after detachment. The opposite directional motors were connected to each other with a no-mass, no-volume cargo in the middle. The load, F, applied on each motor was calculated based on extension (Δl) and stiffness (k_stiff_) of motor as follows,

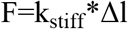

The sign of load applied on each motor is defined by direction of motor stepping. For kinesin, a plus-end-directional transporter, the assisting loads were positive, and the hindering loads were negative. For DDB, a minus-end-directional motor, the assisting loads were negative, and the hindering loads were positive. The model of kinesin’s load-dependent forward stepping rate, kforward was calculated by experimental unloaded velocity (V^0^), stall force (F_stall_), and given constant backward stepping rate (*k_back_*) with following equation,

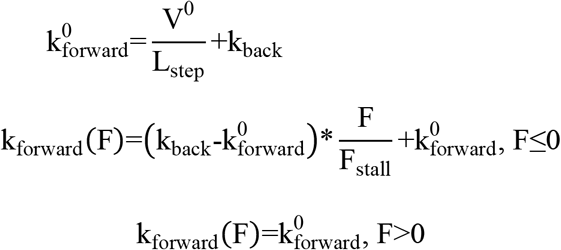

Here, 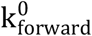 is unloaded forward stepping rate, L_step_ is step size (8 nm), and F is the load applied on motor. With experimental unloaded velocity (V^0^) and run length (RL^0^), the kinesin’s detachment rate, k_detach_, under load was determined by Bell’s model,

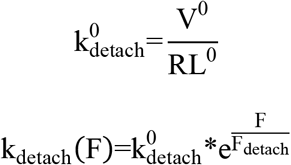

Here, 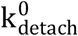 is unloaded detachment rate and F_detach_ is detaching force. For the DDB stepping rate, the basic model followed the force-velocity relationship in Elshenawy et al. (Elshenawy et al., 2019), but the stepping rate under hindering load was modified by a linear function to fit both stall force and unloaded velocity in present experiments.

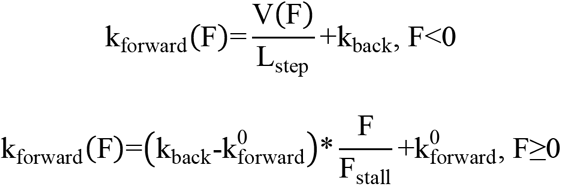

Here, V(F) is the load-dependent velocity in Elshenawy et al. (Elshenawy et al., 2019). The detachment rate of DDB was a given load-independent constant based on previous published simulations.

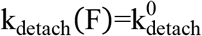

When only one of the motors detached, the position of cargo shifts to where the other motor is bound, and the detached motor can only reattach to the same position with a constant rate. All simulations were run 1000 times and each run was recorded for 50 seconds or until both motors detached from microtubule. All the given parameters are listed in Table 1 in the main text.

For comparing with experimental data, we did data processing on simulation results based on the temporal resolution of tracking experiments. The cargo position was averaged every 285.7 ms and then the instantaneous velocity was calculated for a one second window.

## Supplementary Materials for Gicking et al (2022)

**Figure 1 Supplement 1:**
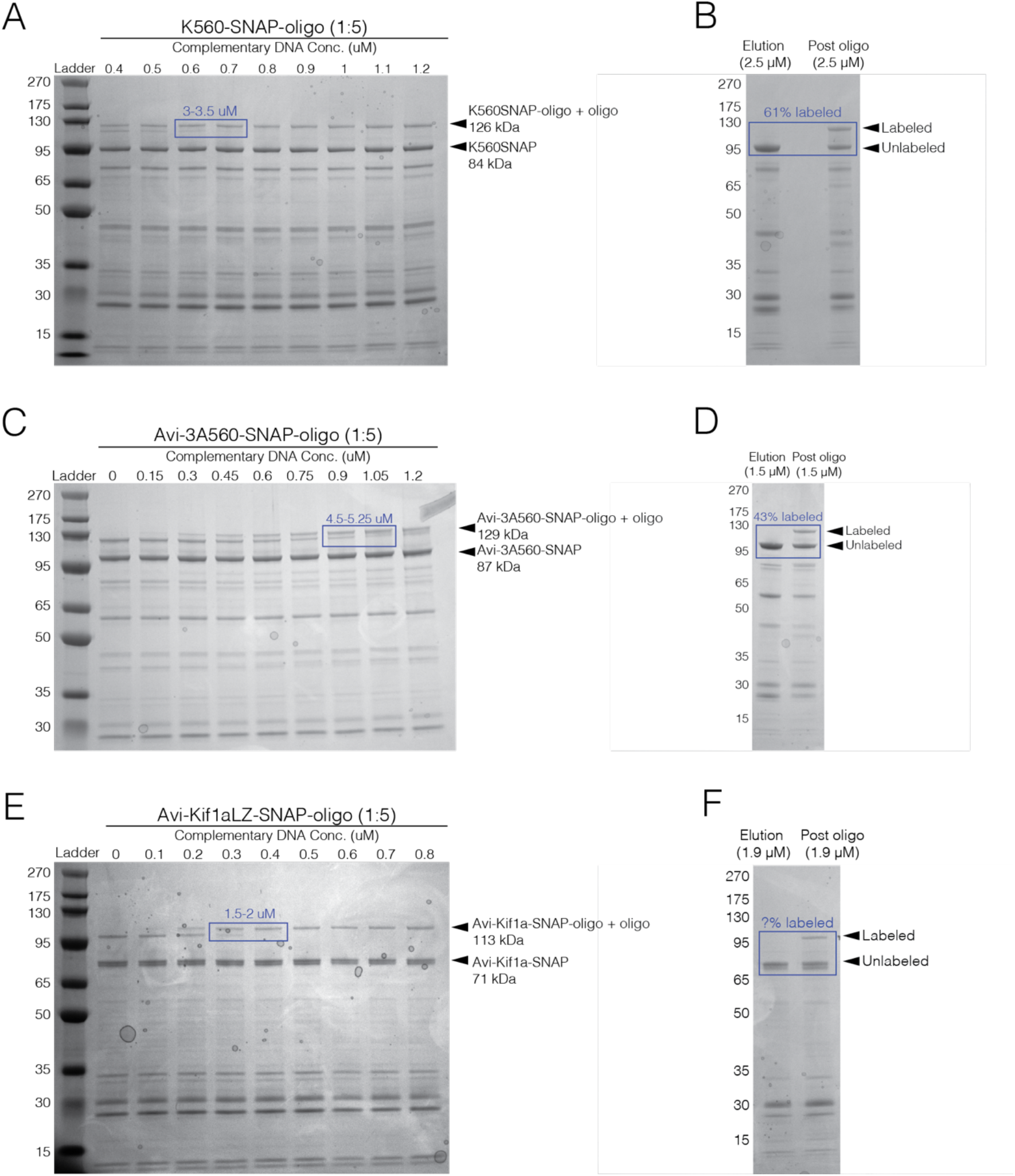
Purification gels and shift assays. (A) Shift assay for K560SNAP, showing the oligo-labeled monomer concentration is 3.25 μM. (B) SDS-PAGE for the purification of K560SNAP showing 63% oligo-labeled monomer. (C) Shift assay for 3A560SNAP, showing oligo-labeled monomer concentration is 4.88 μM. (D) SDS-PAGE for the purification of 3A560SNAP showing 43% oligo-labeled monomer (E) Shift assay for Kif1aLZSNAP, showing effective labeled concentration is 1.75 μM. (F) SDS-PAGE for the purification of Kif1aLZSNAP. The oligo-labeled monomer concentration could not be clearly determined (see methods).

**Figure 1 Supplement 2:**
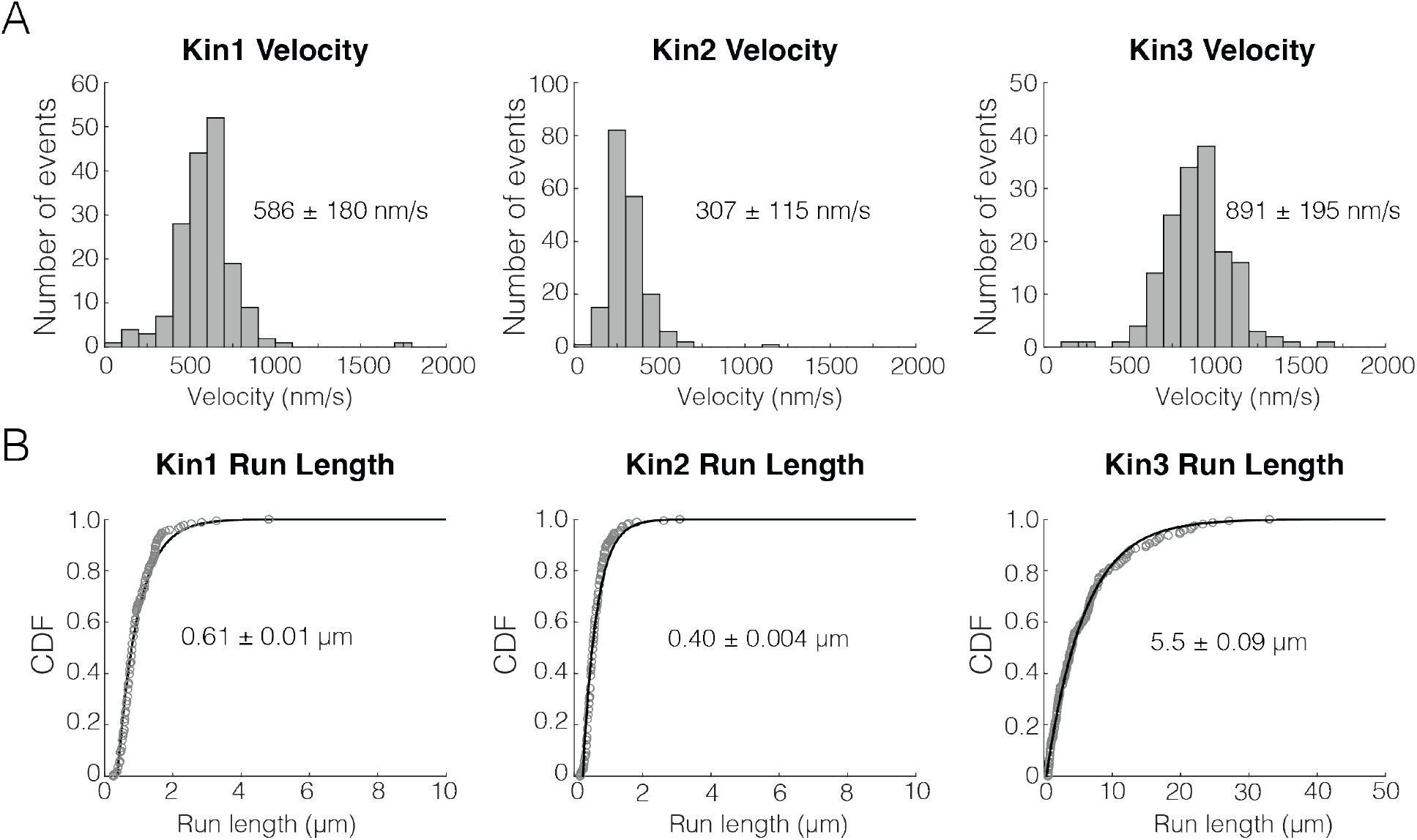
Unloaded run length and velocity for Kin1/2/3. (A) Velocity distributions for kin1, kin2 and kin3. Values represent mean and standard deviation. (B) Cumulative distributions of the run length for Kin1, Kin2 and Kin3. Values represent mean of a single exponential fit and the 95% confidence interval of the bootstrap distribution.

**Figure 1 Supplement 3:**
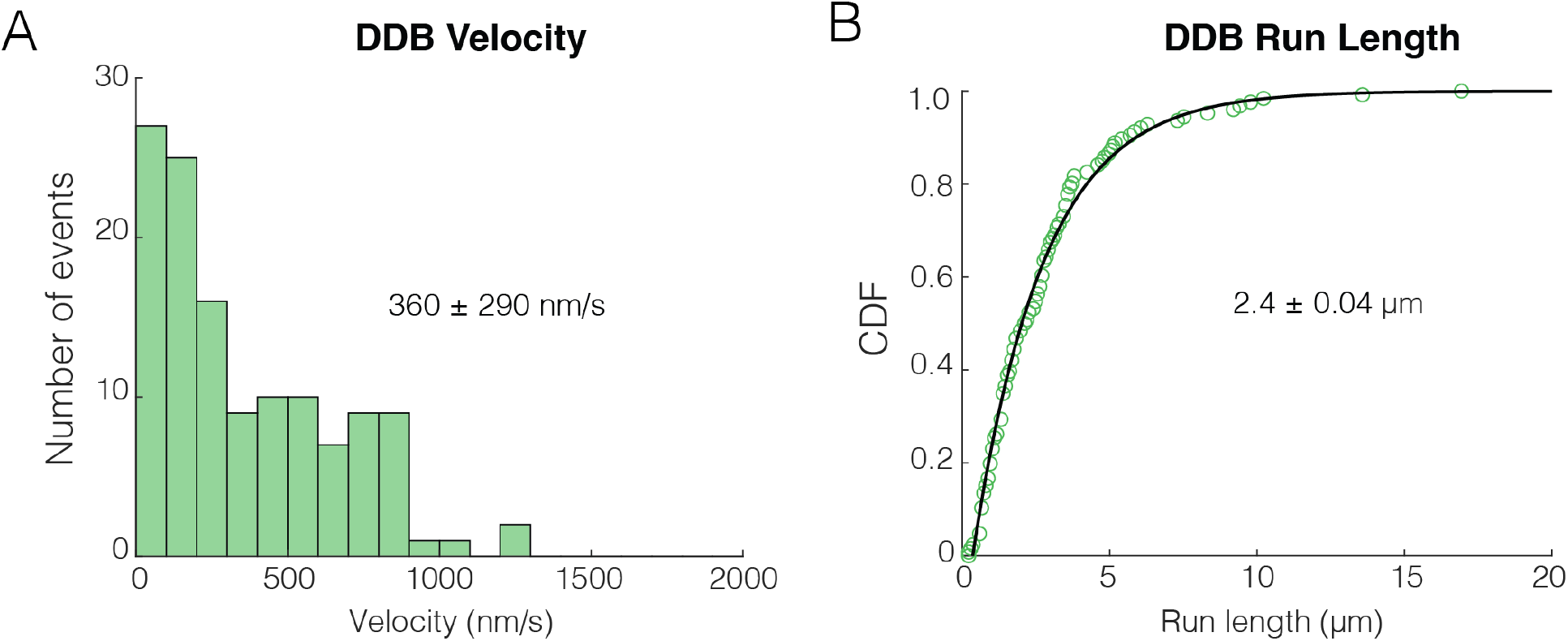
Unloaded run length and velocity for DDB. (A) Velocity distributions for DDB alone. Values represent mean and standard deviation. (B) Cumulative distributions of the run length for DDB alone. Values represent mean of a single exponential fit and the 95% confidence interval of the bootstrap distribution.

**Figure 1 Supplement 4:**
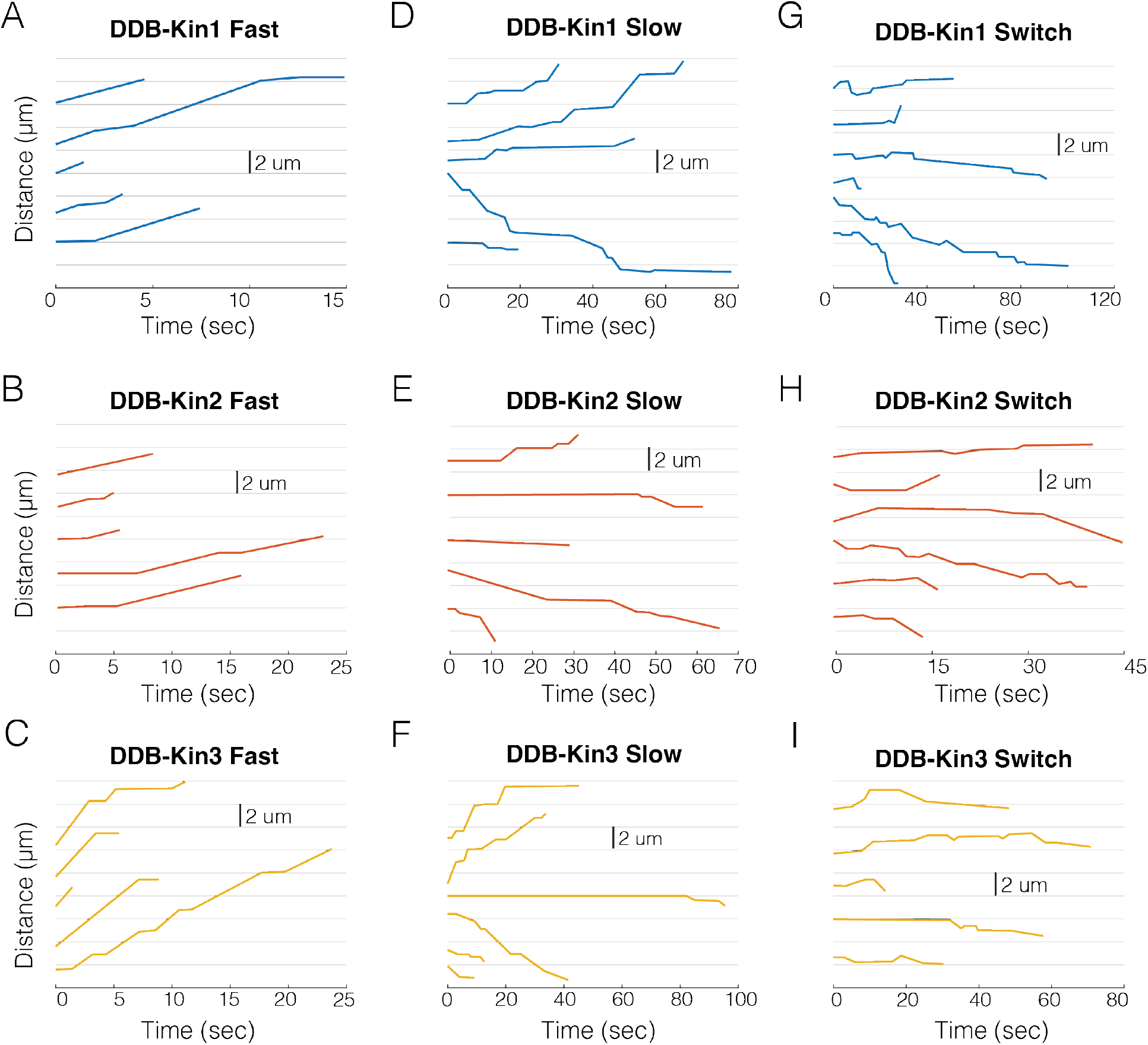
Sample traces for DDB-kin1/2/3 pairs. (A) Sample fast (v > vel threshold) traces for DDB-Kin1 pairs. (B) Sample fast (v > vel threshold) traces for DDB-Kin2 pairs. (C) Sample fast (v > vel threshold) traces for DDB-Kin3 pairs. (D) Sample slow (v < vel threshold) traces for DDB-Kin1 pairs. (E) Sample slow (v < vel threshold) traces for DDB-Kin2 pairs. (F) Sample slow (v < vel threshold) traces for DDB-Kin3 pairs. (G) Examples of traces with directional switching for DDB-Kin1. (H) Examples of traces with directional switching for DDB-Kin2. (I) Examples of traces with directional switching for DDB-Kin3. *Note: time scales are not the same for all plots.*

**Figure 4 Supplement 1:**
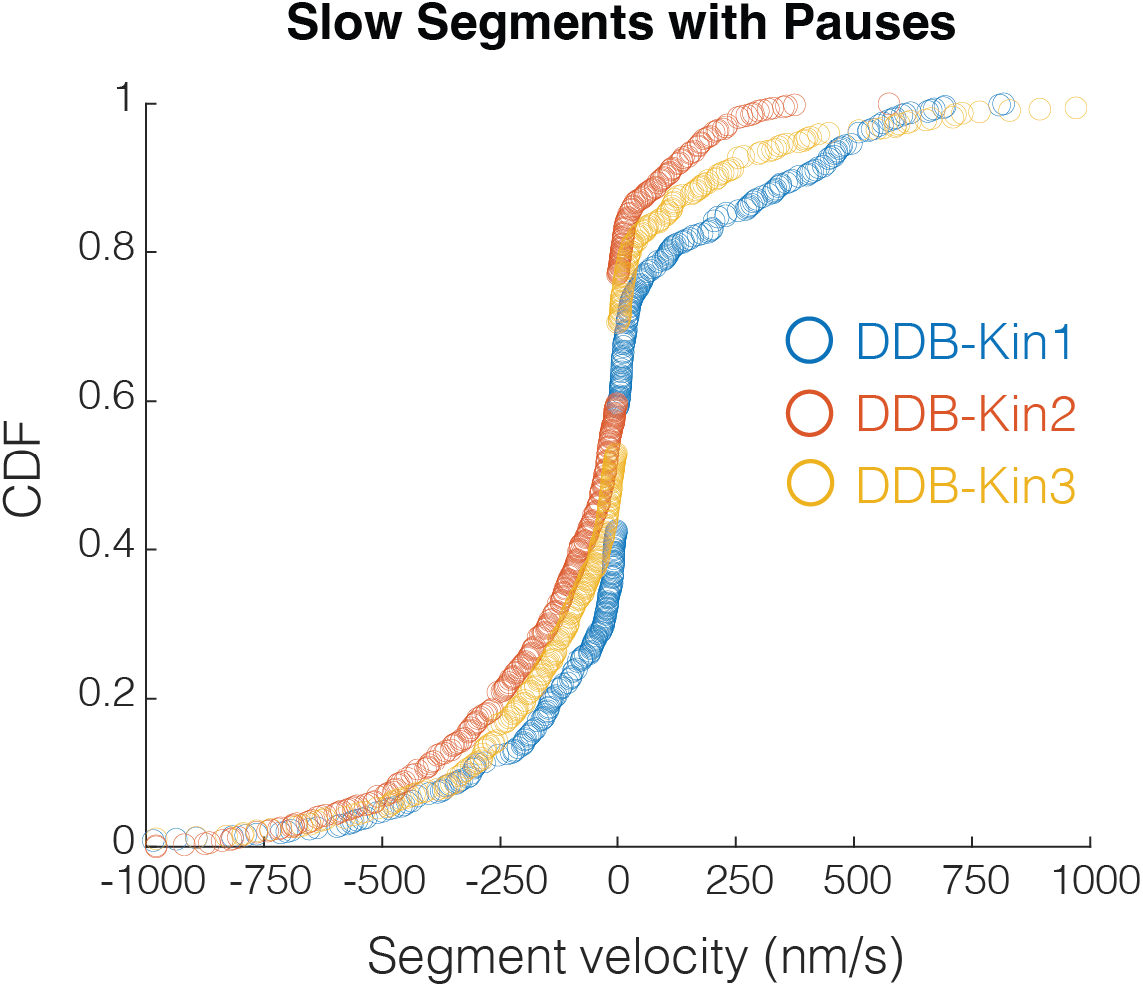
Slow segment velocity distributions. Cumulative distributions of the slow segment (v < threshold) velocities for all three motor pairs.

**Figure 4 Supplement 2:**
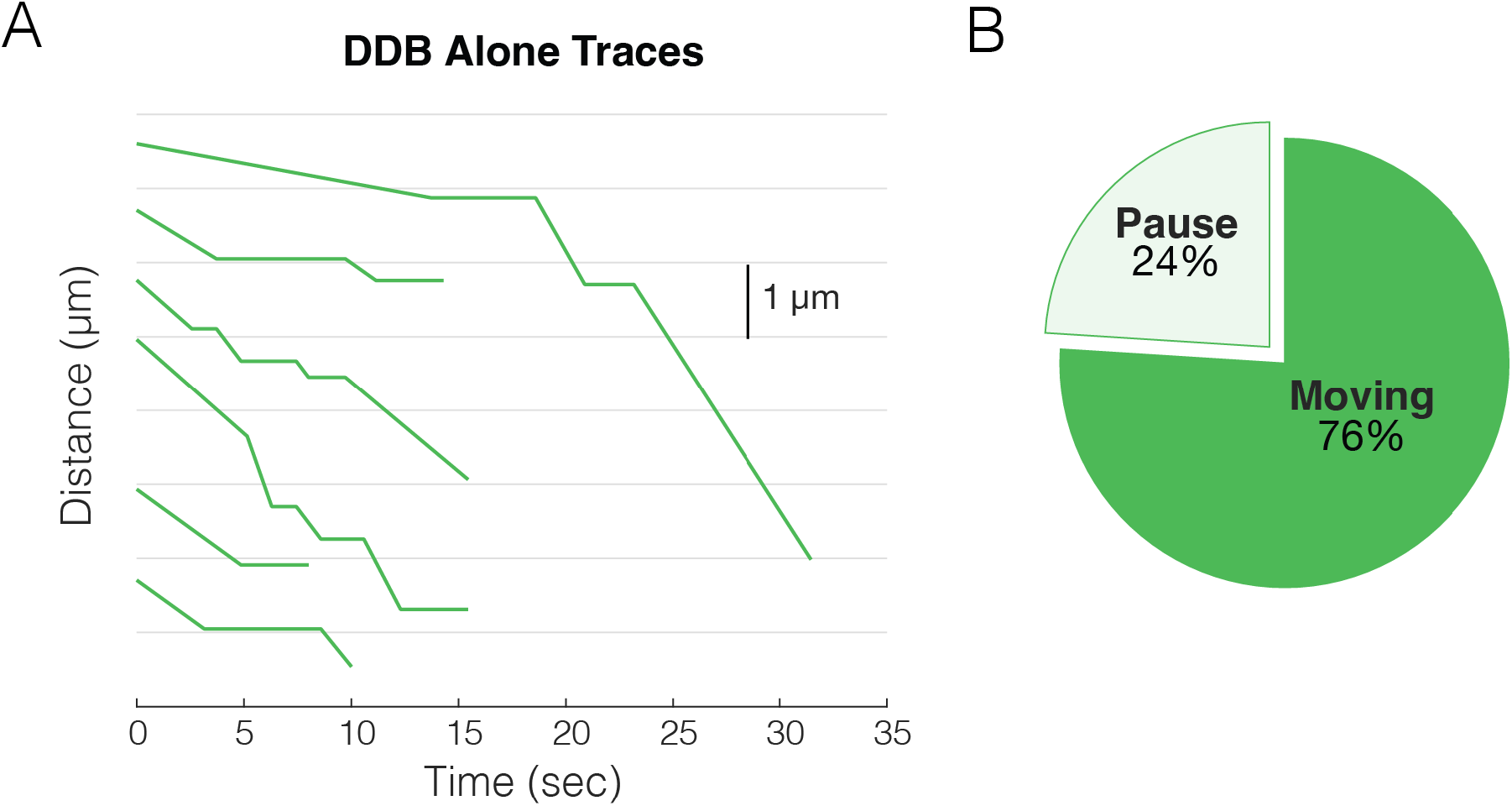
Sample traces for DDB alone. (A) Sample x-t plots for DDB alone. (B) Fraction of paused and moving segments for DDB alone.

**Figure 6 Supplement 1:**
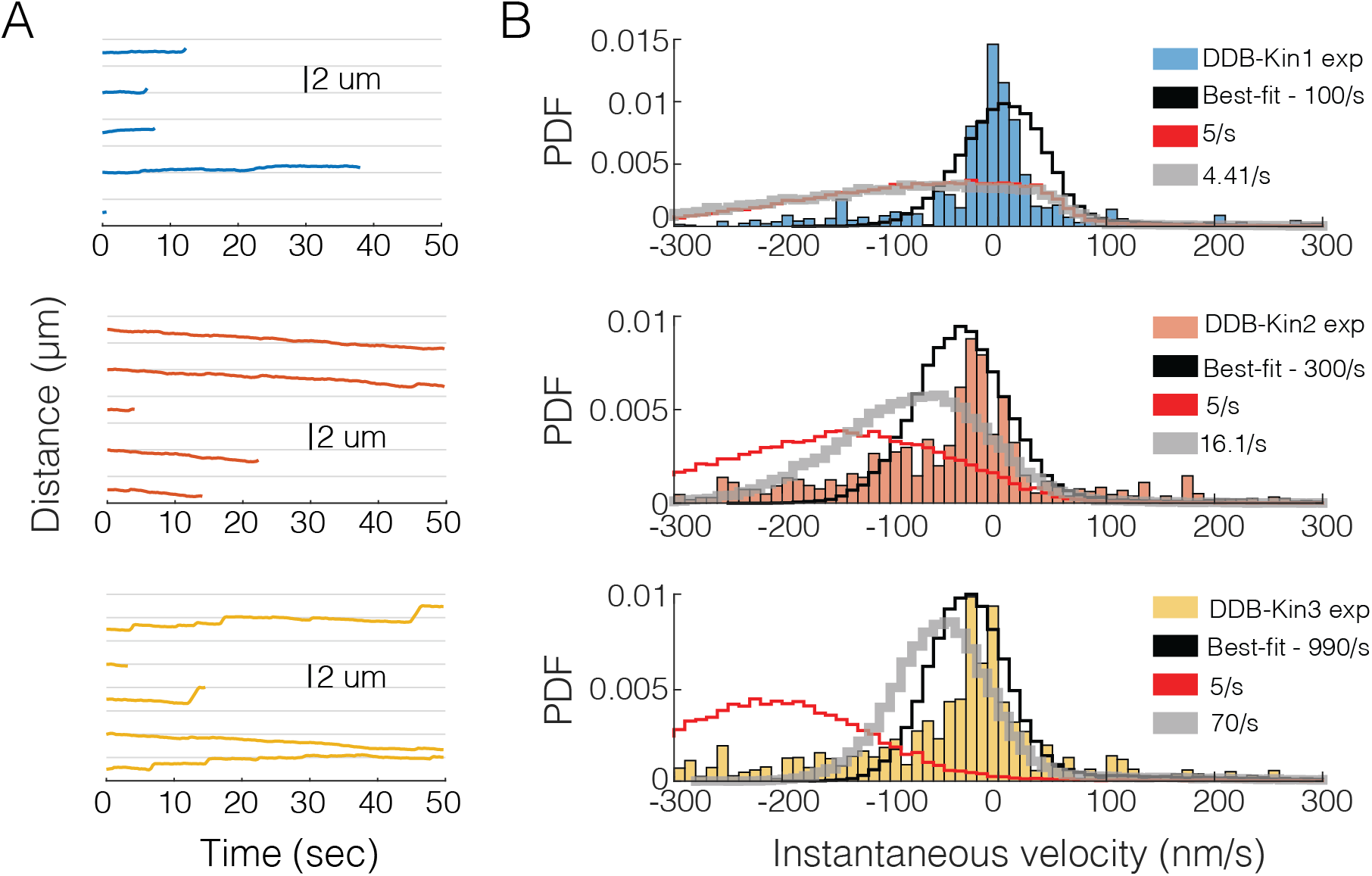
Simulation results for slower reattachment rates. (A) Raw simulation traces using the best-fit model for DDB-Kin1 (top/blue), DDB-Kin2 (middle/orange) and DDB-Kin3 (bottom/yellow). (B) Simulation velocity distributions for DDB-Kin1 (top), DDB-Kin2 (middle), and DDB-Kin3 (bottom) for slower reattachment rates compared with the experimental data (blue/orange/yellow). Best fit model (black) uses parameters listed in Table 1. 5 s^-1^ model (red) uses identical reattachment rate of 5 s^-1^ for each motor. Variable attachment rate model (gray) uses kinesin reattachment rates that are scaled by their bimolecular on-rates for microtubule binding determined by stopped flow (Feng et al., 2018; Zaniewski et al., 2020).

